# KRASG12D mutant cells are outcompeted by wild type neighbours in adult pancreas in an EPHA2-dependent manner

**DOI:** 10.1101/2020.08.04.231050

**Authors:** William Hill, Andreas Zaragkoulias, Beatriz Salvador, Geraint Parfitt, Markella Alatsatianos, Ana Padilha, Sean Porazinski, Thomas E. Woolley, Jennifer P. Morton, Owen J. Sansom, Catherine Hogan

**Affiliations:** European Cancer Stem Cell Research Institute, School of Biosciences, Cardiff University, Hadyn Ellis Building, Maindy road, Cardiff, CF24 4HQ, UK; School of Optometry & Vision Sciences, Cardiff University, Maindy Rd, Cardiff, UK; Faculty of Medicine, St Vincent’s Clinical School, University of NSW, Sydney, Australia; School of Mathematics, Cardiff University, Senghennydd Rd, Cardiff, UK; CRUK Beatson Institute, Glasgow, G61 1BD, UK; Institute of Cancer Sciences, University of Glasgow, G61 1QH, UK

## Abstract

As we age, our tissues are repeatedly challenged by mutational insult, yet cancer occurrence is a relatively rare event. Cells carrying cancer-causing genetic mutations compete with normal neighbours for space and survival in tissues. However, the mechanisms underlying mutant-normal competition in adult tissues and the relevance of this process to cancer remain incompletely understood. Here, we investigate how the adult pancreas maintains tissue health *in vivo* following sporadic expression of oncogenic *Kras* (*KrasG12D*), the key driver mutation in human pancreatic cancer. We find that when present in tissues in low numbers, KrasG12D mutant cells are outcompeted and cleared from exocrine and endocrine compartments *in vivo*. Using quantitative 3D tissue imaging, we show that prior to being cleared, KrasG12D cells lose cell volume, segregate from normal cells and decrease E-cadherin-based cell-cell adhesions with normal neighbours. We identify EphA2 receptor is an essential signal in the clearance of KrasG12D cells from exocrine and endocrine tissues *in vivo*. In the absence of functional EphA2, KrasG12D cells no longer segregate, E-cadherin-based cell-cell adhesions increase and KrasG12D cells are retained in tissues. Retention of KRasG12D cells leads to an increased burden of premalignant pancreatic intraepithelial neoplasia (PanINs) in tissues. Our data show that adult pancreas tissues remodel to clear KrasG12D cells and maintain tissue health. This study provides evidence to support a conserved functional role of EphA2 in Ras-driven cell competition in epithelial tissues and suggests that EphA2 is a novel tumour suppressor in pancreatic cancer.

## Introduction

Epithelial homeostasis is fundamental to survival and is required to balance the number and fitness of cells that contribute to tissue function. Homeostasis is maintained through distinct processes that dynamically maintain this equilibrium in response to tissue crowding^1^, damage^2^ or mutational insult^3–10^. Retention of excess, mutant or aberrant cells would impair tissue integrity and promote disease^11–13^. The mechanisms underlying these processes are multifaceted, and involve cell competition^14,15^, mechanical cues^16,17^ and cell plasticity^18^. Epithelial cells expressing oncogenes compete for space and survival in tissues and are often eliminated via processes that require the presence of normal cells; however, the mechanisms underlying how normal cells sense and eliminate mutant cells remain incompletely understood. We recently identified differential EphA2 signalling as a novel, evolutionary conserved mechanism that drives the segregation and elimination of RasV12 cells in simple epithelia^10,19^. EphA2 is a receptor tyrosine kinase of the Eph-ephrin family of cell-cell communication signals^20^ that play a general role in regulating cell proliferation and survival, and cell-cell adhesion at tissue boundaries leading to compartmentalisation of cells^21^. Whether EphA2 is a general regulator of mammalian tissue homeostasis *in vivo* is currently unknown.

The pancreas is composed of two functionally distinct compartments of epithelial cells derived from common progenitors^22^. Exocrine acinar cells produce and secrete digestive enzymes that travel to the gut via ductal networks. Endocrine cells of the islets of Langerhans regulate blood glucose by producing and secreting hormones such as insulin and glucagon. Pancreatic cancer arises from activating mutations in oncogenic Kras^23^ and the majority (90%) of human pancreatic tumours develop sporadically from cells carrying *KRAS* mutations^24^; however, KRasG12D mutations alone are insufficient to drive malignancy^25,26^. Pancreatic ductal adenocarcinoma (PDAC; the most common form of human pancreatic cancer) develops predominantly from pancreatic intraepithelial neoplasia (PanIN)^27^. The cancer cell of origin in PDAC remains controversial; however, *in vivo* mouse studies indicate that tumorigenesis can develop from cells of exocrine and endocrine lineages^*28–30*^. Unlike rapidly proliferating epithelia such as intestine or skin, the adult pancreas is not actively renewing, has limited proliferative capacity during homeostasis and relies on cell plasticity to regenerate in response to injury^31,32^. What is less understood is how the adult pancreas maintains tissue health following mutational insult. Here, we set out to address this question and investigate the requirement of EphA2 in adult pancreas following sporadic, sparse induction of KrasG12D expressing cells *in vivo*, recapitulating a scenario of sporadic tumorigenesis. Using fluorescence imaging of murine pancreas tissues and quantitative image analysis platforms, we demonstrate that KrasG12D cells are actively cleared from adult pancreas over time and in an EphA2-dependent manner. In the absence of functional EphA2 receptor, mutant cells are retained and the number of premalignant lesions in tissues increases, suggesting that EphA2-driven competition is tumour suppressive.

## Results

### Sparse KrasG12D mutant cells are lost from adult pancreas tissues over time

Epithelial cells expressing oncogenic Ras (HrasV12) are often eliminated from normal epithelial tissues and this is regarded to be tumour preventative^4,6,10,11^. Since *KRAS* mutations are key driver mutations of human pancreatic cancer, we set out to determine whether cells expressing oncogenic Kras (KrasG12D) also compete for space and survival in pancreas tissues *in vivo*. To address this, we used the pancreas-specific *Pdx1-Cre^ER^ LSL-Kras^G12D/+^; Rosa26^LSL-RFP^* (KC) mouse^26,33^. The KC mouse accurately models human PanIN development and spontaneous progression to invasive PDAC^33^ and is clinically relevant to human disease. Experimental controls were *Pdx1-Cre^ERT^, Rosa26^LSL-RFP^* (*Kras* wild type; Control). To generate tissues mosaic for *Kras* expression, we administered a single low dose of tamoxifen to 6-8 week old mice and induced *Pdx1*-Cre recombinase in a low number of cells in an otherwise normal epithelium. Genetic recombination also induced expression of red fluorescent protein (RFP) reporter in *Kras* cells. We found that low dose tamoxifen induced stochastic RFP labelling of both endocrine and exocrine lineages in adult tissues at low frequency, with RFP positive acinar and islet cells predominantly represented (Fig. 1A). To determine the fate of RFP-labelled cells, we monitored global RFP fluorescence in fresh frozen tissues over time and found that a single low dose of tamoxifen induced RFP labelling in approximately 20% of the tissue (SFig. 1A). When comparing tissues harvested at 7 days post induction (p.i.) of Cre recombinase, we found comparable levels of RFP in KrasG12D and *Kras* wild type control tissues (Fig. 1B, left panels; 1C, round data points, p=0.07), indicating that initial levels of recombination were consistent between the two experimental groups. We also observed minimal variation in the levels of recombination when sampling from the tail, middle or head of the pancreas (SFig. 1B). In contrast, administration of a high dose of tamoxifen^28^, resulted in RFP expression in approximately 80% of the tissue (SFig. 1A).

**Figure 1:**
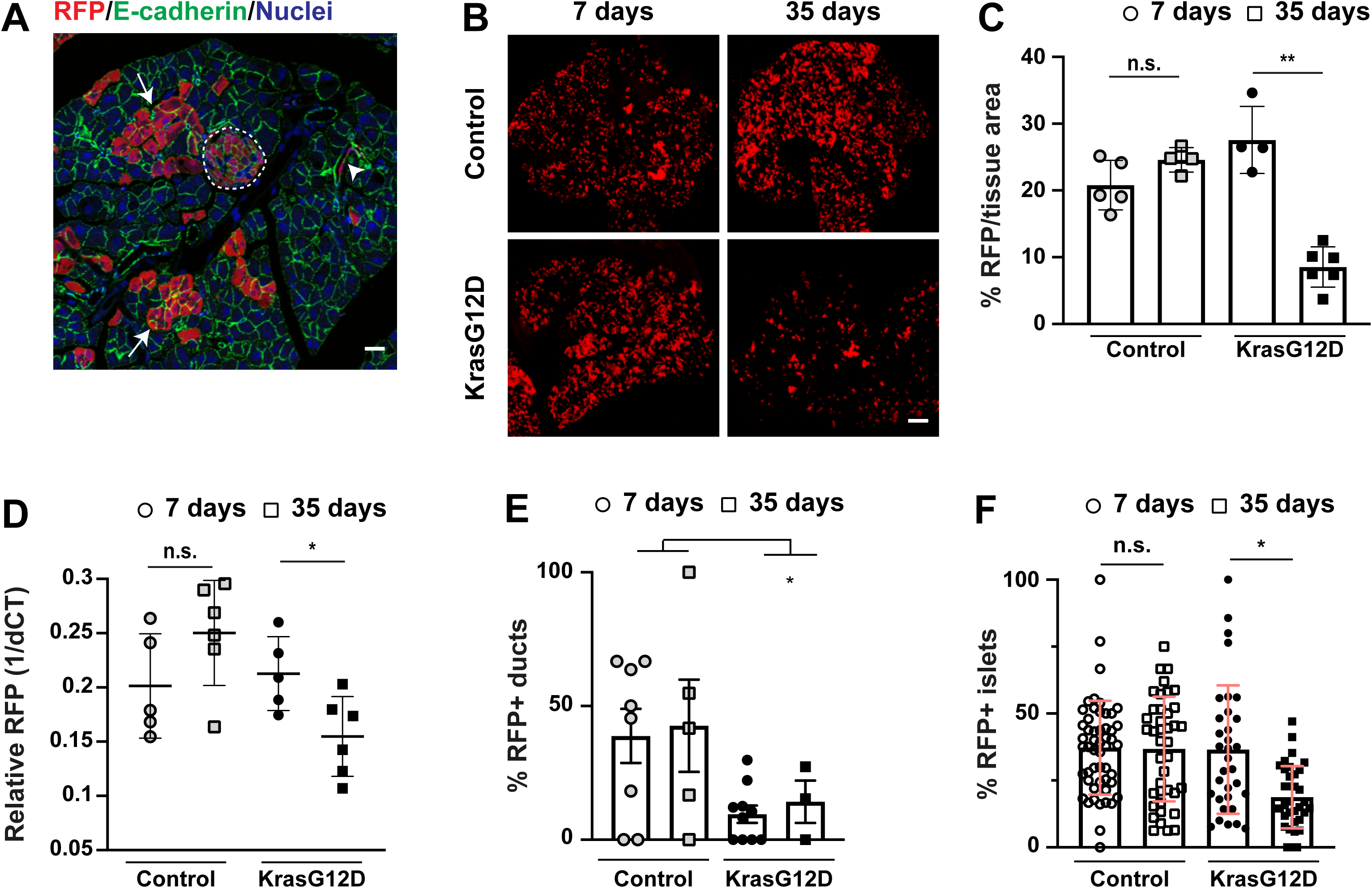
RFP-labelled KrasG12D cells are cleared from exocrine and endocrine compartments *in vivo*. (**A**) Representative confocal image of murine pancreas tissue at 7 days post induction (p.i.) of Cre recombinase. Fixed tissue was processed using immunofluorescence tomography protocols, and stained with anti-RFP (red), anti-E-cadherin (green) antibodies and Hoescht (blue). White arrows: RFP labelled acinar cells; white arrowhead: RFP labelled ductal epithelial cells; dashed white line: RFP labelled islet. (**B**) Representative stitched confocal tile scan images of fresh frozen murine pancreas tissues showing endogenous RFP fluorescence. Tissues were harvested from *Kras* wild type control (top) or KrasG12D (bottom) animals at 7 days (left panels) or 35 days (right panels) post induction (p.i.). (**C**) Bar graph showing percentage endogenous RFP fluorescence per tissue area in *Kras* wild type control (grey points) or KrasG12D (black points) tissues, harvested at 7-days (circles) or 35-days (squares) p.i. Data are mean +/− s.d. of RFP fluorescence averaged from five tissue slices (of 50 µm apart) per mouse. n.s.=not significant. **p=0.002, unpaired Student t test using Welch correction; n=5 *Kras* wild type controls at 7 and 35 days, n=4 and n=6 for KrasG12D at 7 days and 35 days, respectively. (**D**) Scatter plot of RFP expression in genomic DNA relative to housekeeping gene in tissues harvested at 7 days (circles) and 35 days (squares) from *Kras* wild type control (grey points) and KrasG12D (black points) mice. Data are mean +/− s.d. of RFP expression/tissue per mouse. n.s.=not significant. *p=0.024, unpaired Student t test using Welch correction; n=5 and n=6 for *Kras* wild type controls at 7 days and 35 days, respectively; n=5 and n=6 for KrasG12D at 7 days and 35 days, respectively. (**E**) Bar graph of percentage of RFP positive ducts in *Kras* wild type control and KrasG12D tissues over time. Data are mean +/− s.e.m positive ducts. *p=0.04, one-way ANOVA; n=8 and n=5 mice for *Kras* wild type controls at 7 days (grey circles) and 35 days (white squares), respectively; n=10 and n=3 mice for KrasG12D at 7 days (black circles) and 35 days (black squares), respectively. (**F**) Percentage RFP positive islet cells/total islet cells in *Kras* wild type control and KrasG12D tissues at 7 days (circles) and 35 days (squares) p.i. Data represent mean +/− s.d. islets pooled from n=4 mice/genotype. n.s.=not significant. ***p=0.0008, non-parametric Student t test. n=52 and n=40 islets for *Kras* wild type controls at 7 days (grey circles) and 35 days (grey squares), respectively; n=33 and 23 n=35 islets for KrasG12D at 7 days (black circles) and 35 days (black squares), respectively. Scale bars, 500 µm.

Next, we induced recombination in cohorts of *KrasG12D* (KC) and *Kras* wild type animals with single low dose tamoxifen and harvested tissue at specific time points (days p.i.) to monitor *Kras* cell fate. We have previously shown using whole-body imaging experiments that GFP-tagged KrasG12D cells are selectively lost during the first four weeks of life and early pancreas development^26^. Therefore, we reasoned that putative competition between KrasG12D and normal cells in adult tissues would occur over protracted time points. To assess this, we chose 35 days p.i. as an end point and monitored the amount of RFP fluorescence per tissue area in fresh frozen tissues over time. Taking this approach, we found that the proportion of RFP positive tissue did not significantly change in control animals between 7 and 35 days p.i. (Fig. 1B, upper panels; p=0.092; 1C, grey points). In contrast, RFP fluorescence significantly decreased over time in KC tissues (Fig 1B, lower panels; p=0.002; 1C, black points), suggesting that KrasG12D cells are cleared from adult tissues over a four-week period. To validate these findings at the genetic level, we analysed relative levels of recombined RFP in genomic DNA isolated from pancreas. Consistent with global RFP fluorescence data, we found that the relative amount of recombined RFP in genomic DNA from the pancreas significantly decreased in KC tissues (p=0.024; Fig. 1D, black points), whereas levels of recombined RFP remained constant in *Kras* wild type controls (p=0.13; Fig. 1D, grey points). Crucially and in contrast to that observed in developing tissues^26^, the proportion of RFP positive tissue remained unchanged over time in adult KC tissues treated with high dose tamoxifen (SFig. 1C, lower panels; p=0.5; 1D, black points), suggesting that selective loss of RFP positive cells from adult KrasG12D tissues is not cell autonomous and requires the presence of normal cells. Consistent with previous reports^11^, we found that RFP positive ducts were significantly less frequent in KC tissues compared to *Kras* wild type controls (p=0.04; Fig 1E). Similarly, the number of RFP positive islet cells significantly decreased over time in KrasG12D tissues only (p=0.0008; Fig 1F). Since exocrine acinar cells are predominately labelled with RFP (Fig. 1A), we conclude that KrasG12D cells are outcompeted by normal cells in exocrine and endocrine epithelial compartments in adult pancreas *in vivo*.

### KrasG12D mutant cells are outcompeted in an EphA2-dependent manner

To determine the molecular mechanisms underlying KrasG12D cell competition, we first monitored tissue homeostasis by scoring cell proliferation and cell death events in fixed tissues. Using TUNEL assays (SFig. 2A) and immunostaining for cleaved caspase 3 (SFig. 2B), we found that apoptosis events were extremely rare in both KrasG12D and *Kras* wild type tissues; the very low number of cleaved caspase 3 positive events observed remained unchanged over time regardless of genotype (p>0.1; SFig. 2B), suggesting that clearance of KrasG12D cells is unlikely to require apoptosis. We also found no evidence of cell senescence in tissues (not shown). Interestingly, we found that the number of Ki67 positive cells per tissue area significantly decreased over time in both KrasG12D and *Kras* wild type tissues (p=0.003; SFig. 2C), suggesting that cell proliferation decreases in aging pancreas tissues over time^34,35^. Our data also imply that changes in cell proliferation rates are unlikely to contribute to clearance of KrasG12D cells from KC tissues.

**Figure 2:**
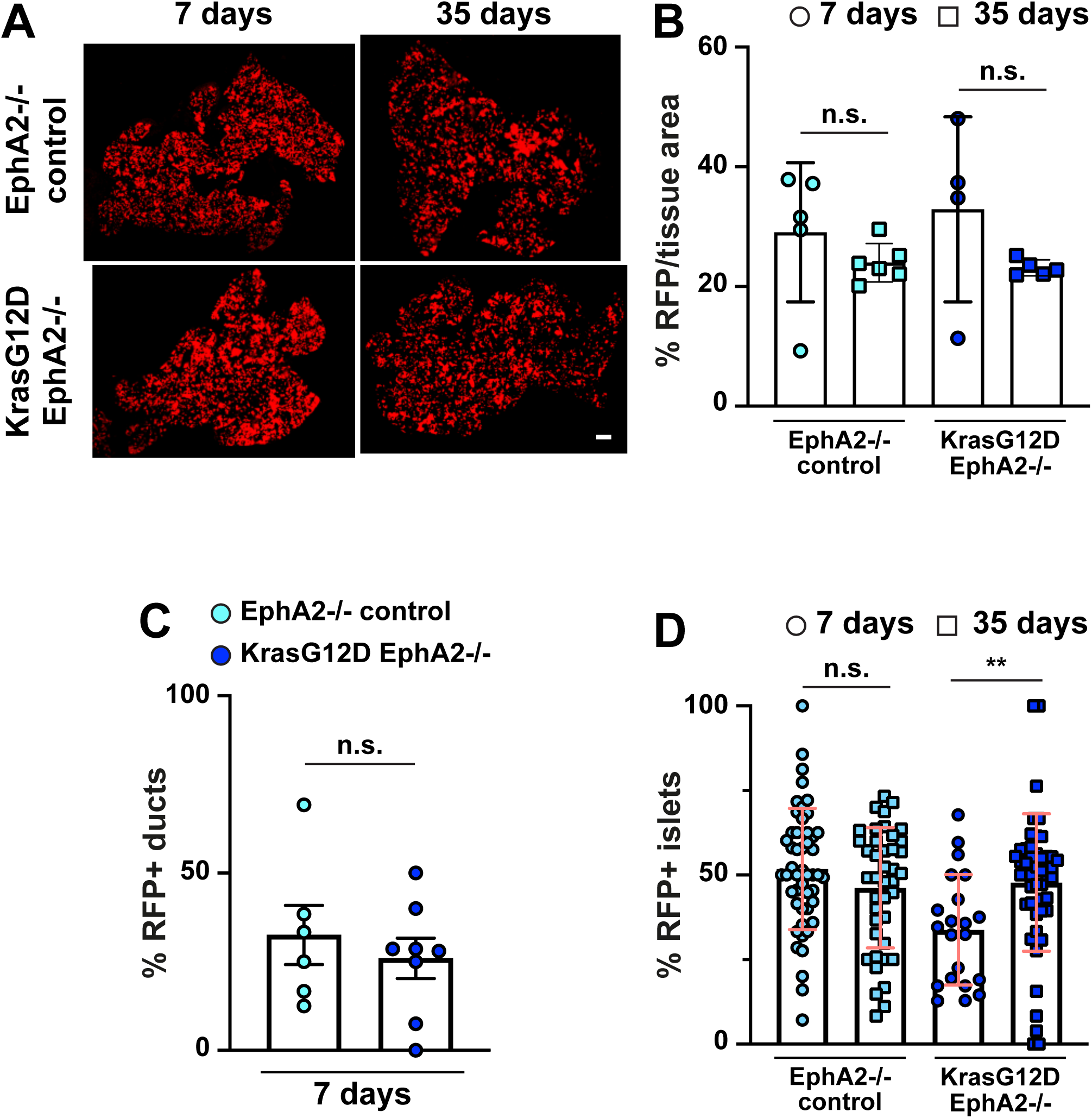
KrasG12D cells are retained in EphA2 knockout tissues. (**A**) Representative stitched confocal tile scan images of fresh frozen murine pancreas tissues showing endogenous RFP fluorescence. Tissues were harvested from EphA2 knockout (EphA2−/− control; top) or KrasG12D/+ EphA2−/− (bottom) animals at 7 days (left panels) or 35 days (right panels) p.i. (**B**) Bar graph showing percentage endogenous RFP fluorescence per tissue area in EphA2−/− control or KrasG12D EphA2−/− tissues, harvested at 7 days (circles) or 35 days (squares) p.i. Data are mean +/− s.d. of RFP fluorescence averaged from five tissue slices (of 50 µm apart) per mouse. n.s.=not significant. n=5 and n=6 mice for EphA2−/− controls at 7 days (light blue circles) and 35 days (light blue squares), respectively. n=4 and n=5 mice for KrasG12D EphA2−/− at 7 days (blue circles) and 35 days (blue squares), respectively (**C**) Bar graph of proportion of exocrine ducts positive for RFP in EphA2−/− control (light blue circles) and KrasG12D EphA2−/− (blue circles) tissues at 7 days p.i. Data are mean +/− s.e.m positive ducts. n.s.=not significant. n=6 EphA2−/− control and n=8 KrasG12D EphA2−/− mice. (**D**) Percentage RFP positive islet cells/total islet cells in EphA2−/− control and KrasG12D EphA2−/− tissues at 7 days (circles) and 35 days (squares) p.i. Data represent mean +/− s.d. islets pooled from n=4 mice/genotype at 7 days; n=6 mice/genotype at 35 days. **p=0.0035, non-parametric Student t test. n=49 and n=41 islets for EphA2−/− controls at 7 days (light blue circles) and 35 days (light blue squares), respectively; n=21 and n=47 islets for KrasG12D EphA2−/− at 7 days (blue circles) and 35 days (blue squares), respectively. Scale bar, 500 µm.

Based on our previous work describing a role of EphA2 in the elimination RasV12 cells from simple epithelia^10,19^, we asked whether EphA2 is required to promote the clearance of KrasG12D cells from adult pancreas tissues *in vivo*. To test this, we crossed KC animals onto EphA2 knockout mice, generating *Pdx1*-Cre^ERT^; *LSL-Kras*^G12D^; *Rosa26*^LSL-RFP^; *EphA2*^−/−^ mice (referred to as KCE). *Pdx1*-Cre^ERT^; *Rosa26*^LSL-RFP^; *EphA2*^−/−^ (*EphA2*^−/−^ control) animals were included as experimental controls. Mice homozygous for the targeted EphA2 mutation are deficient for EphA2 protein^36^ and are therefore knockout for EphA2. Comparing levels of RFP fluorescence in fresh frozen tissues harvested at 7 days p.i. indicated no significant difference in the level of recombination between different cohorts (p=0.15, EphA2−/− versus *Kras* wild type control; p=0.34, KrasG12D EphA2−/− versus KrasG12D). Similar to *Kras* wild type control tissues, we observed that the proportion of RFP positive tissue did not significantly change in *EphA2*−/− control tissues over time (Fig. 2A, top panels; p=0.57; 2B, light blue points). Notably and in contrast to KrasG12D tissues (Fig. 1B–D), RFP fluorescence was not significantly different over time in KrasG12D EphA2−/− tissues (Fig. 2A, lower panels; p=0.29; 2B, KrasG12D EphA2−/−: blue points), indicating that RFP positive KrasG12D cells are not cleared from tissues depleted of EphA2. Moreover, the number of RFP positive ducts present in tissues at 7 days p.i. were similar in KrasG12D EphA2−/− and EphA2−/− control tissues (p=0.53, Fig. 2C), and were significantly increased in KrasG12D EphA2−/− tissues compared to KrasG12D tissues (p=0.028). Similarly, RFP positive islets significantly increased in KrasG12D EphA2−/− tissues over time (p=0.0035; Fig. 2D). We observed no significant difference in the number of cleaved caspase-3 positive cells over time in KrasG12D EphA2−/− tissues (p=0.18; SFig. 2D, blue points). Cell proliferation events significantly decreased in aging KrasG12D EphA2−/− tissues and EphA2 −/− controls (p=0.04; SFig. 2E), similar to that observed in KrasG12D or Kras wild type tissues. We conclude that KrasG12D cells are outcompeted from both exocrine and endocrine compartments in an EphA2-dependent manner, and independent of changes in global cell proliferation/cell death. Moreover, KrasG12D cells are retained in EphA2 depleted tissues.

### KrasG12D cells segregate in an EphA2-dependent manner

Competitive interactions with normal cells induce RasV12 cells to adopt a highly contractile morphology and segregate from normal neighbours^4,10,19^. To gain insight into KrasG12D cell fate *in vivo*, we examined KrasG12D cell morphology in fixed pancreas tissues using immunofluorescence tomography (IT)^37,38^ and quantitative image analysis. Fixed, serial tissue sections were immunolabelled for E-cadherin and RFP before being imaged by confocal microscopy. Next, we aligned and stacked serial tissue sections to generate 3D reconstructions (SMovie 1) and individual cells were segmented based on E-cadherin staining. Using inter-nuclear distance measurements between cells in direct contact^10^, we found that RFP positive acinar cells formed significantly more compact clusters in KC tissues compared to *Kras* wild type controls (p<0.0001; Fig. 3A). We also quantified cell volume and found that RFP positive acinar cells significantly decreased in cell volume in KC tissues compared to *Kras* wild type controls (p<0.0001; Fig. 3B). Together, these data suggest that KrasG12D acinar cells shrink and segregate from normal cells *in vivo*. Next, we examined the morphology of RFP positive acinar cells in *EphA2* knockout tissues at 7 days p.i. We found that RFP positive KrasG12D cells no longer formed tightly packed clusters in KrasG12D EphA2−/− tissues (p<0.0001; Fig. 3A). In addition, cell volume of RFP positive KrasG12D cells was significantly increased in KrasG12D EphA2−/− tissues compared to KrasG12D tissues (p=0.003; Fig.3B). Together, our data indicate that EphA2 is required to promote the segregation of KrasG12D acinar cells from normal neighbours *in vivo*.

**Figure 3:**
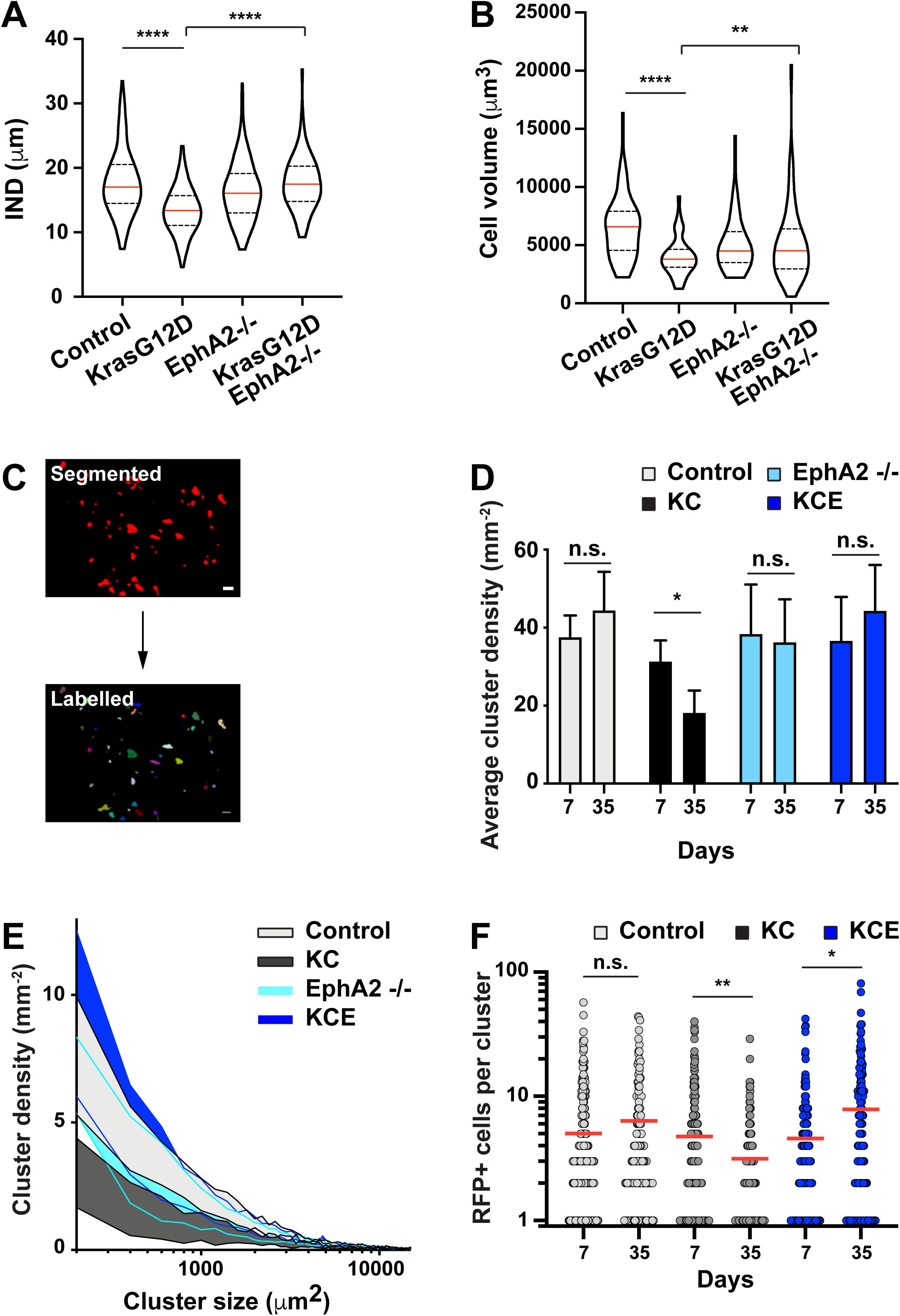
KrasG12D cells segregate in tissues *in vivo* in an EphA2-dependent manner and are eliminated as single cells. Violin plots of (**A**) inter nuclear distance (IND, µm) between neighbouring RFP positive cells and (**B**) RFP positive cell volume (µm^3^) in tissues harvested at 7 days p.i. Red line: median; dashed lines: quartiles. Data represent values pooled from 3 mice (*Kras* wild type controls, KrasG12D EphA2−/−, EphA2−/− controls) or 4 mice (KrasG12D). (A) n=184 cells *Kras* wild type control; n=218 cells KrasG12D; n=162 cells KrasG12D EphA2−/−; n=78 24 cells EphA2−/− controls (B) n=87 cells *Kras* wild type control; n=89 cells KrasG12D; n=113 cells KrasG12D EphA2−/−; n=60 cells EphA2−/− controls. ****p<0.0001, **p=0.0031, non-parametric Student t tests in (A) and unpaired Student t tests using Welch correction in (B). (**C**) Representative images of segmented RFP fluorescence, which labels individual clusters (pseudo-colour). Scale bar, 500 µm. (**D**) Bar graph of average cluster density per tissue area from five tissue sections (of 50 µm apart) per mouse. Cluster density was quantified from tissues harvested at 7 days and 35 days p.i. Light grey bar: *Kras* wild type control; Black bar: KrasG12D (KC); Light blue bar: EphA2−/− control; Blue bar: KrasG12D, EphA2−/− (KCE). n.s.=not significant; *p=0.025, unpaired Student t test using Welch correction. (**E**) Frequency distribution graph of clusters of varying size (mm-2). Density of small clusters (<2000 µm^2^ in size) is lower in KrasG12D tissues (KC; dark grey curve) at 35 days p.i., compared to *Kras* wild type controls (light grey curve), EphA2−/− controls (light blue curve) and KrasG12D EphA2−/− (KCE; blue curve). For (D) and (E) n=5 mice/17892 clusters (7 days) and n=4 mice/12967 clusters (35 days) *Kras* wild type control; n=3 mice/15263 clusters (7 days) n=6 mice/6089 clusters (35 days) KrasG12D (KC); n=5 mice/16942 clusters (7 days) n=6/23305 clusters (35 days) EphA2−/− control; n=4 mice/13422 clusters (7 days) n=5 mice/18402 clusters (35 days) KrasG12D EphA2−/− (KCE). (**F**) Scatter plot of number of RFP positive acinar cells per cluster in tissues harvested at 7 days or 35 days p.i. Light grey: Kras wild type control; Dark grey: KrasG12D (KC); Blue: KrasG12D, EphA2−/− (KCE). Red bar denotes mean. Data represent values pooled from 3 mice (*Kras* wild type controls, KrasG12D EphA2−/−, EphA2−/− controls) or 4 mice (KrasG12D). n.s.=not significant; *p=0.017, **p=0.002, non-parametric Student t tests. Control: n=312 (7 days) and n=136 (35 days); KrasG12D (KC): n=252 (7 days) and n=143 (35 days); KrasG12D EphA2−/− (KCE): n=212 (7 days) and 186 (35 days).

To further investigate the requirement of EphA2 in driving these phenotypes, we isolated primary murine pancreatic ductal epithelial cells (PDEC) and applied established coculture assays *in vitro*^4,10^. When mixed with non-transformed PDECs at 1:50 ratios, pre-labelled KrasG12D ductal epithelial cells formed tightly packed clusters (SFig. 3A; KR:N, top panels), and significantly decreased in cell area in a non-autonomous manner (p<0.0001; SFig. 3B, KR:N versus KR:KR). Moreover, quantification of the index of sphericity indicated that clusters of KrasG12D cells were significantly more round when surrounded by normal cells (KR:N) compared to KR:KR controls (p<0.0001; SFig. 3C), suggesting that KrasG12D cells separate from normal neighbours *in vitro* via the formation of smooth and distinct boundaries. Crucially, KrasG12D cells depleted for EphA2 (KRE cells) no longer adopted a contractile phenotype (SFig 3A; KRE:N, middle panels), had a significantly higher cluster area when surrounded by normal cells (p<0.0001; SFig 3B, KR:N compared to KRE:N), and segregated less efficiently from normal neighbours compared to KrasG12D cells (p<0.0001; SFig. 3C, KR:N versus KRE:N). Notably, we also observed apical extrusion of KrasG12D cells from normal monolayers in an EphA2-dependent manner (SFig 3D, white arrows). These data suggest that EphA2 expressed on KrasG12D cells is required to promote cell segregation and extrusion of KrasG12D pancreatic ductal epithelial cells *in vitro*. Taken together, our data show that exocrine KrasG12D cells become contractile and segregate from normal cells in an EphA2-dependent manner.

### KrasG12D cells are outcompeted at normal-mutant boundaries

Our data provide strong evidence that competitive interactions with normal cells induce KrasG12D cells to segregate *in vivo*. Based on our previous observations in MDCK cells^10^, we speculated that KrasG12D cells are eliminated at normal-mutant cell-cell boundaries *in vivo*. To test this hypothesis, we developed a stochastic mathematical model of mutant-normal cell competition. In the model, mutant cells in direct contact with normal cells are outcompeted (SFig. 4A). Simulations of clusters of mutant cells of varying sizes reveals that mutant cells are eliminated from the periphery of a mutant cluster, irrespective of cluster size, leading to an overall decrease in observed cluster size (SFig. 4A, bottom panels). The model also suggests that small clusters of mutant cells would be lost first (SFig. 4B). To examine this experimentally, we segmented global RFP fluorescence data to quantify RFP-labelled clusters in tissues (Fig. 3C) and determine cluster density and the distribution of clusters based on size. We found that cluster density (p=0.28; Fig. 3D, grey bars) and the distribution of clusters of varying size (SFig. 4C) did not significantly change over time in *Kras* wild type tissues. In contrast, cluster density significantly decreased over time in KrasG12D (KC) tissues (p=0.025; Fig. 3D, black bars). We also observed significantly fewer smaller clusters (of <2000 μm^2^) at 35 days p.i. in KrasG12D tissues compared to *Kras* wild type controls (Fig. 3E and SFig. 4D, black line; SFig. 4G, WT v KC), supporting the model that mutant cells are more likely to be eliminated when present in tissues in low numbers. Consistent with global RFP data (Fig. 2B), cluster density (p=0.35; Fig. 3D, blue bars) and the distribution of clusters based on size did not significantly change in EphA2−/− control (SFig. 4E) or in KrasG12D EphA2−/− (SFig. 4F) tissues over time. Moreover, we scored significantly more small clusters in KrasG12D EphA2−/− (KCE) tissues at 35 days p.i. compared to KrasG12D (KC) tissues at the same time point (SFig. 3G, KC v KCE), with size distribution of RFP positive clusters in KrasG12D EphA2−/− (KCE) tissues comparable to controls (Fig. 3E), further supporting an essential role of EphA2 in KrasG12D cell competition. To control for the observed changes in acinar cell morphology *in vivo* (Fig. 3A, 3B), we quantified the number of RFP positive acinar cells per cluster at the single cell level. This revealed that the number of RFP positive cells per cluster significantly decreased in KrasG12D (KC) tissues over time and not *Kras* wild type controls, irrespective of cluster size (p=0.002; Fig 3F, dark grey points). In contrast, RFP positive cells per cluster significantly increased in KrasG12D EphA2−/− (KCE) tissues (p=0.017; Fig. 3F, blue points). These data support the model and suggest that direct competition with normal cells leads to a progressive loss of mutant cells and an overall decrease in cluster size, in an EphA2 dependent manner.

**Figure 4:**
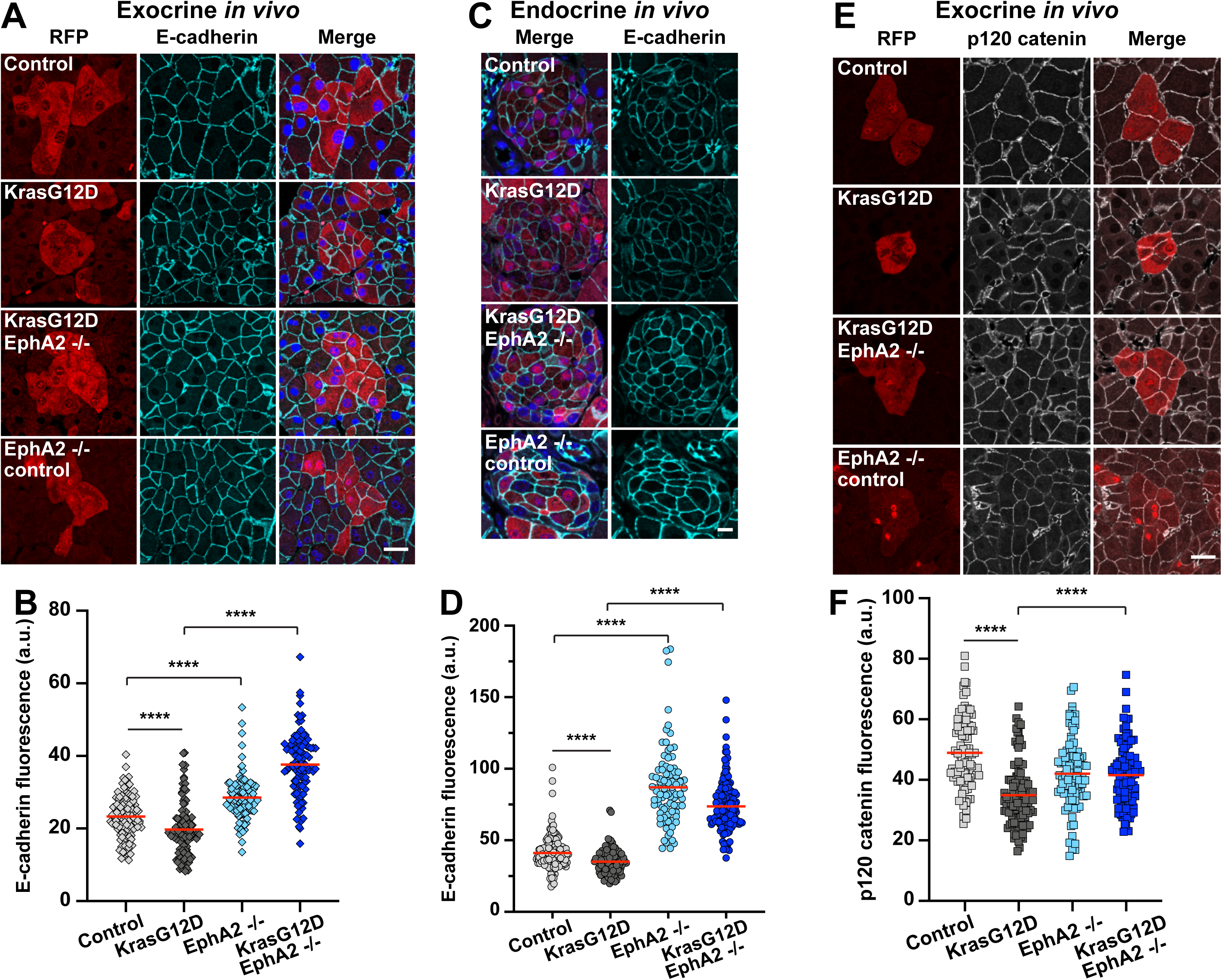
Adherens junctions are remodelled at mutant-normal cell-cell contacts in exocrine and endocrine tissues and in an EphA2-dependent manner. (**A**) Confocal images of exocrine acinar cells in tissues from each genotype fixed at 7 days p.i. Tissues were processed using immunofluorescence tomography protocols and stained with anti-RFP (red), anti-E-cadherin (cyan) antibodies and Hoescht (blue). (**B**) Scatter plot of E-cadherin fluorescence at cell-cell interface 25 between RFP positive and RFP negative cells in *Kras* wild type control (light grey), KrasG12D (dark grey), EphA2−/− (light blue); KrasG12D, EphA2−/− (blue) tissues. Red bar denotes mean. Data represent cell-cell contacts pooled from n=3 mice (*Kras* wild type controls, KrasG12D EphA2−/−, EphA2−/− controls), n=4 mice (KrasG12D). ****p<0.0001, non-parametric Student t tests. n=111 Control; n=106 KrasG12D; n=98 EphA2−/− control; n=96 KrasG12D EphA2−/−. (**C**) Confocal images of endocrine islets in tissues from each genotype fixed at 7 days p.i. Tissues were processed using immunofluorescence tomography protocols and stained with anti-RFP (red), anti-E-cadherin (cyan) antibodies and Hoescht (blue). (**D**) Scatter plot of E-cadherin fluorescence at cell-cell interface between RFP positive and RFP negative cells in *Kras* wild type control (light grey), KrasG12D (dark grey), EphA2−/− (light blue); KrasG12D, EphA2−/− (blue) tissues. Red bar denotes mean. Data represent cell-cell contacts pooled from n=3 mice (*Kras* wild type control, EphA2−/− control, KrasG12D EphA2−/−), n=4 mice KrasG12D. ****p<0.0001, non-parametric Student t tests. n=152 Control; n=100 KrasG12D; n=90 EphA2−/− control; n=138 KrasG12D EphA2−/−. (**E**) Confocal images of exocrine acinar cells in tissues from each genotype fixed at 7 days p.i. Tissues were processed using immunofluorescence tomography protocols and stained with anti-RFP (red), anti-p120 catenin (grey) antibodies. (**F**) Scatter plot of p120 catenin fluorescence at cell-cell interface between RFP positive and RFP negative cells in *Kras* wild type control (light grey), KrasG12D (dark grey), EphA2−/− (light blue); KrasG12D EphA2−/− (blue) tissues. Red bar denotes mean. Data represent cell-cell contacts pooled from 3 mice/genotype. ****p<0.0001, nonparametric Student t tests. n=92 Control; n=111 KrasG12D; n=96 EphA2−/− control; n=93 KrasG12D EphA2−/−. Scale bars, 20 µm.

### EphA2 is required to destabilise E-cadherin-based cell-cell contacts at normal-mutant boundaries

We have previously shown that RasV12 cells segregate from normal cells in an E-cadherin-dependent manner^10^. We reasoned that E-cadherin-based cell-cell contacts between mutant and normal cells must be remodelled *in vivo* in response to a loss of mutant cell volume and prior to mutant cell elimination. Moreover, EphA2 may play a pivotal role in this process by triggering cell repulsion at mutant-normal cell boundaries^10^. To investigate this, we immunostained fixed tissues harvested at 7 days p.i. for endogenous E-cadherin. We focused on cell-cell contacts between RFP positive cells and unlabelled neighbours in exocrine acinar and endocrine compartments. We found that E-cadherin appeared to localise uniformly at cell-cell contacts in *Kras* wild-type tissues (Fig. 4A and 4C, Control, upper panels) but appeared weaker and punctate at cell-cell contacts in KC tissues (Fig. 4A and 4C, KrasG12D panels). To confirm this observation, we quantified E-cadherin fluorescence at cell-cell contacts between RFP positive and RFP negative cells. We found that E-cadherin was significantly decreased at RFP positive/RFP negative cell-cell interfaces in exocrine acinar (p<0.0001; Fig. 4B) and endocrine islet (p<0.0001; Fig 4D) compartments in KC tissues compared to *Kras* wild-type controls. Similarly, we found that E-cadherin was predominantly intracellular in KrasG12D ductal epithelial cells in a non-cell autonomous manner *in vitro* (SFig. 3E; KR:N). In contrast, E-cadherin was significantly enriched at cell-cell contacts between RFP positive cells and RFP negative neighbours in exocrine and endocrine KCE tissues *in vivo* (Fig. 4A and 4C; KrasG12D EphA2−/− panels; p<0.0001; Fig. 4B, D). We also observed that E-cadherin was no longer intracellular in KRE cells surrounded by non-transformed cells *in vitro* but appeared uniformly localised at cell-cell contacts (SFig. 3E; KRE:N panels). Together, our data suggest that E-cadherin localisation at normal-mutant cell-cell contacts is dependent on EphA2. Interestingly, E-cadherin fluorescence was significantly higher at RFP positive/RFP negative boundaries in exocrine and endocrine *EphA2*−/− control tissues compared to *Kras* wild type controls (p<0.0001; Fig. 4B, 4D), suggesting that EphA2 regulates E-cadherin localisation at cell-cell contacts in a cell autonomous manner.

We speculated that EphA2 may promote E-cadherin turnover at mutant-normal cellcell contacts. With this in mind, we stained tissues fixed at 7 days p.i. for endogenous p120 catenin, which directly binds to the intracellular domain of Ecadherin and stabilises cell-cell adhesion by masking motifs necessary for engagement of the endocytic machinery^39,40^. Focusing on exocrine acinar cells, we found that p120 catenin poorly localised to cell-cell contacts between RFP positive and unlabelled cells in KC tissues compared to *Kras* wild type controls (Fig. 4E). Quantification of p120 catenin fluorescence at cell-cell contacts demonstrated that levels of p120 catenin were significantly lower at RFP positive/RFP negative cell-cell contacts in KC tissues compared to *Kras* wild type control (p<0.0001; Fig. 4F). In contrast, p120 catenin fluorescence was significantly increased at RFP positive/RFP negative cell-cell contacts in KCE tissue compared to KC tissue (Fig. 4E; p<0.0001; 4F), suggesting that EphA2 is required to regulate E-cadherin turnover at cell-cell contacts in exocrine pancreas *in vivo*. Remarkably, our data also demonstrate that E-cadherin-based cell-cell contacts are remodelled specifically at KrasG12D and normal cell-cells boundaries and this is concomitant with a loss in KrasG12D cell volume.

### Normal cells expand to compensate for the loss of mutant cells

In rapidly proliferating tissues, elimination of aberrant cells is often accompanied by compensatory proliferation of surrounding winner cells to replace lost tissue^41^. Since we observed a significant decrease in cell proliferation over time in all tissues (SFig. 2C, 2E), we postulated that loss of mutant cells from KrasG12D tissues triggers compensatory expansion of adjacent wild type cells rather than cell proliferation. Using 3D reconstructed tissue datasets, we quantified cell volume of normal acinar cells either directly adjacent or non-adjacent to RFP positive acinar cells (Fig. 5A). We found that in KrasG12D (KC) tissues, normal cells directly adjacent to RFP positive cells had a significantly larger cell volume, compared to that of non-adjacent normal cells (p<0.0001; Fig. 5B), suggesting that normal cells in direct contact with KrasG12D cells expand in size. In contrast, we found that in KrasG12D EphA2−/− (KCE) tissues, cell volume of normal cells adjacent to RFP positive cells was not significantly different to that of non-adjacent cells (p=0.46; Fig. 5C), suggesting that expansion of normal cells requires functional EphA2. Considering that RFP positive cells lose cell volume in KrasG12D (KC) tissues and not in KrasG12D EphA2−/− (KCE) tissues (Fig 3B and Fig. 5B, 5C), our data imply that normal cells adjacent to KrasG12D cells expand in size to compensate for the loss of KrasG12D cell volume in an EphA2 dependent manner.

**Figure 5:**
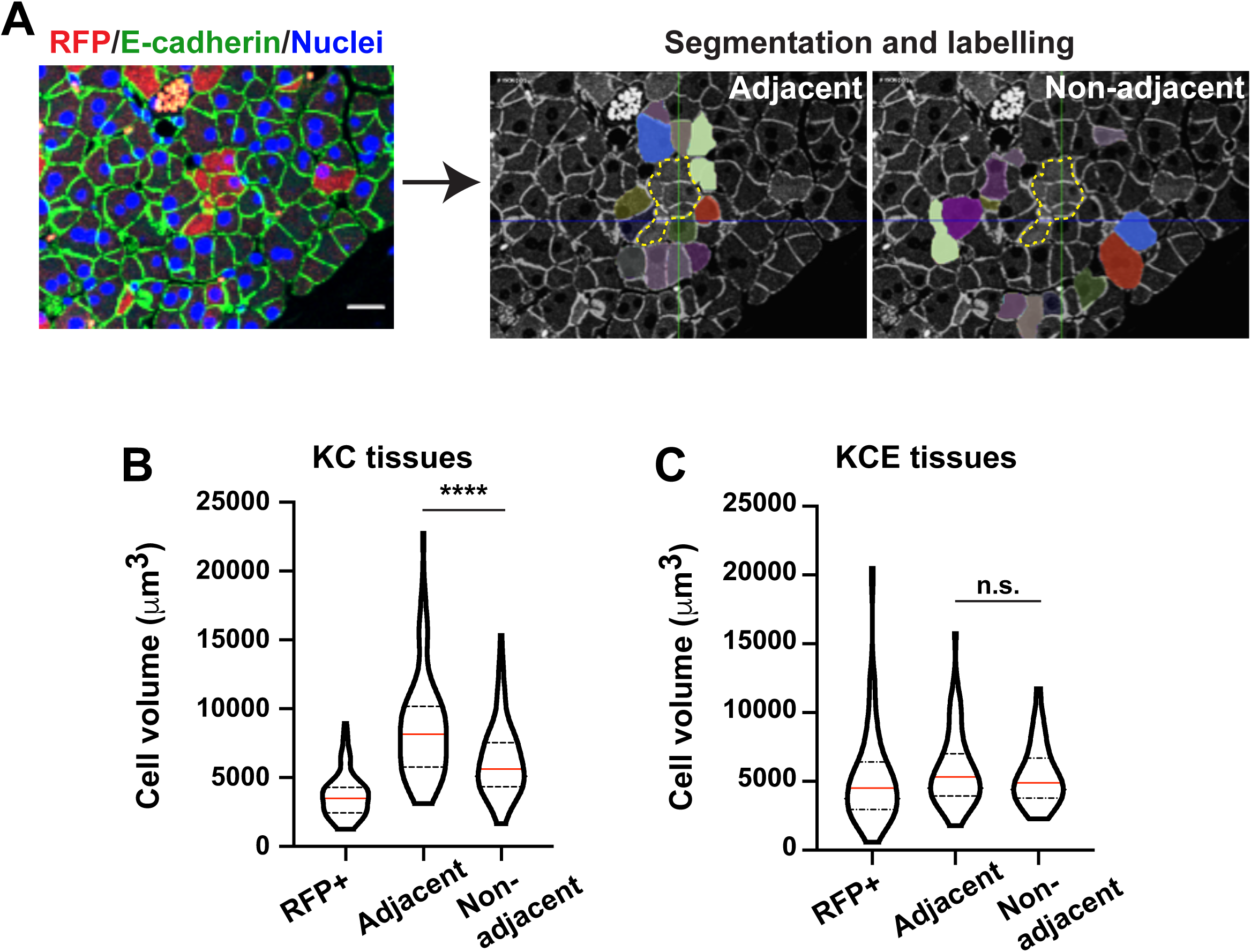
Normal cells directly neighbouring KrasG12D cells increase in cell volume *in vivo* and in an EphA2-dependent manner. (**A**) Confocal image of pancreas tissue fixed at 7 days p.i. and stained with anti-RFP (red) and anti-Ecadherin (green) antibodies and Hoescht (blue) using immunofluorescence tomography protocols. Segmentation analysis in 3D labels normal cells (pseudocoloured) neighbouring RFP positive mutant cells (dashed line). (**B-C**) Violin plots of cell volume (µm3) of RFP positive and unlabelled normal cells, adjacent or nonadjacent to RFP positive cells in (**B**) KrasG12D (KC) and (**C**) KrasG12D EphA2−/− 26 (KCE) tissues. Red line denotes median; dashed lines: quartiles. n.s.=not significant; ****p<0.0001, calculated using non-parametric Student t tests. Data represent volume of individual cells pooled from n=3 mice KrasG12D EphA2−/− (KCE); n=4 mice KrasG12D (KC). KrasG12D (KC) tissues: n=101 RFP positive cells; n=91 adjacent normal; n=84 non-adjacent normal. KrasG12D EphA2−/− (KCE): n=113 RFP positive cells; n=61 adjacent normal; n=61 non-adjacent normal. Scale bar, 20 µm.

### Loss of EphA2 potentiates early development of premalignant lesions

Finally, we asked whether retention of KrasG12D cells in EphA2 knockout tissues promotes development of premalignant lesions. It is generally accepted that expression of KrasG12D in pancreas tissues triggers the development of premalignant pancreatic intraepithelial neoplasia (PanIN); however, PDAC tumour development requires additional genetic mutations^26^ or chronic inflammation^25^. While the majority of KrasG12D cells were eliminated by 35 days p.i., a rare population of KrasG12D cells consistently remained in KC tissues (Fig 1B, C). Using Alcian blue staining as a marker of mucin positive PanIN lesions^42^, we first investigated whether the presence of this rare population of KrasG12D cells in tissues could induce PanIN lesion development. Indeed, we observed that PanIN lesions were rare in KrasG12D (KC) tissues at 35 days p.i. (Fig. 6B; 2/7 mice) but were more frequent at 140 days p.i., (Fig. 6A, B; 7/9 mice), suggesting that non-eliminated KrasG12D cells expand to form lesions over protracted time points. These data are consistent with our previously reported data^26^ and suggest that a subpopulation of KrasG12D cells evade elimination signals to initiate disease. Strikingly, we observed significantly more mucin positive PanIN lesions in KrasG12D EphA2−/− (KCE) tissues at 35 days p.i. compared to KrasG12D (KC) tissues at the same time point (p=0.003; Fig. 6A, B; 6/6 mice), suggesting that loss of EphA2 signalling increases PanIN lesion burden in tissues. We conclude that loss of functional EphA2 leads to an increase in the number of KrasG12D cells retained in tissues, which in turn increases the frequency of oncogenic niches and premalignant lesion development.

**Figure 6:**
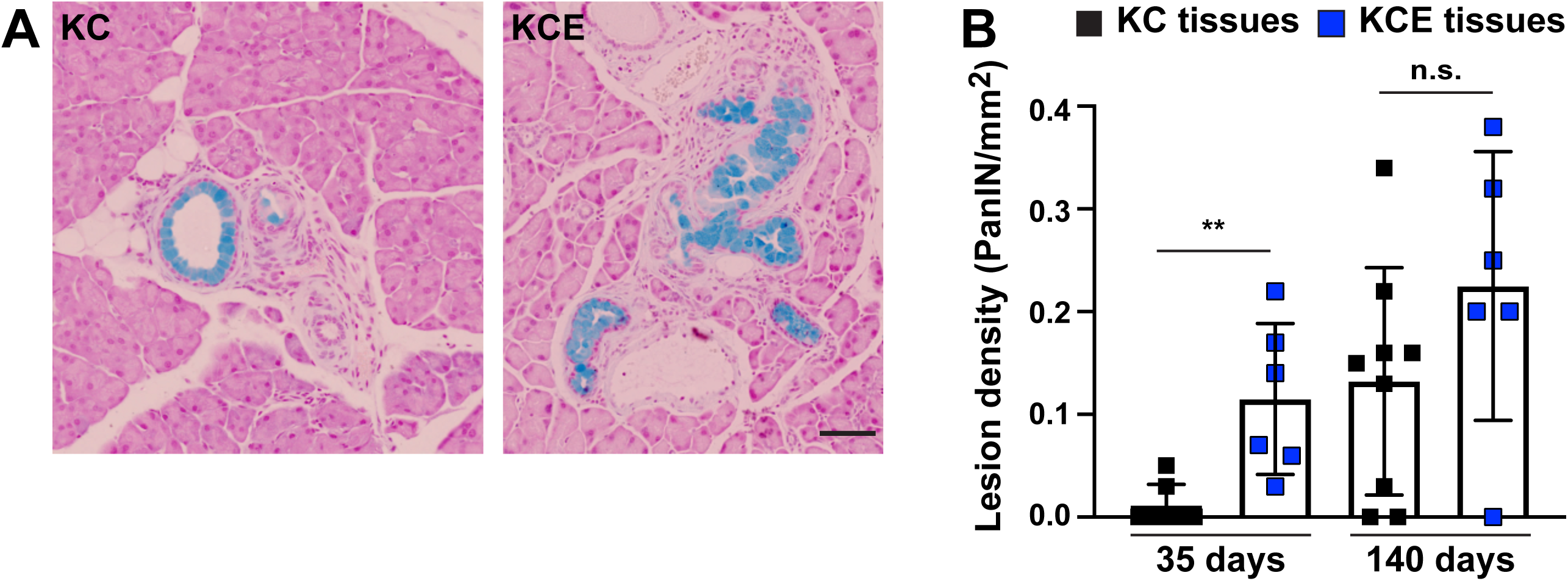
Premalignant PanIN lesions are more abundant in KrasG12D EphA2 knockout tissues. (**A**) Representative bright field images of pancreas tissues fixed at 140 days p.i. and stained for mucin (blue). Scale bar, 100 µm. (**B**) Bar chart of mucin positive premalignant lesion density (PanIN/mm^2^) in KrasG12D (KC; black) or KrasG12D EphA2−/− (KCE; blue) tissues over time. Data represent mean +/− s.d. of average number of lesions per tissue area from four FFPE tissue sections (100 µm apart) per mouse. n.s.=not significant; **p=0.0029, non-parametric Student t test. KrasG12D (KC): n=7 (35 days) and n=9 mice (140 days); KrasG12D EphA2−/− (KCE): n=6 for 35 days and 140 days

## Discussion

In this study, we show that cells carrying oncogenic KrasG12D mutations are outcompeted by normal cells in adult pancreas epithelia *in vivo*. We reveal a novel essential role of EphA2 receptor in this process and show that EphA2 is required to remove KrasG12D cells from both exocrine and endocrine compartments *in vivo*. Loss of EphA2 leads to the retention of KrasG12D cells and the concomitant increase in premalignant PanIN lesion burden in tissues. Thus, our data reveal that EphA2 is required to maintain pancreas tissue health by promoting the elimination of KrasG12D cells, with retention of mutant cells increasing the probability of tumour initiation. Our data add to the growing body of evidence^7,43–45^ demonstrating an innate ability of epithelial tissues to prevent tumour initiation following mutational insult.

Here, we show that competition with normal cells leads to the clearance of KrasG12D cells from pancreas tissues. Our data suggest that elimination of KrasG12D cells from pancreas epithelia occurs over different time scales depending on the local tissue architecture; e.g., single layer of ductal epithelial cells versus a 3D acinus or islet, and that different processes may be required to remove KrasG12D cells from different tissue compartments. In general, cell competition induces apoptosis of unfit cells and cell proliferation in surrounding cells, thus ensuring overall tissue size is maintained^41^. We found no evidence of apoptosis of KrasG12D cells *in vivo*, suggesting that mutant cells are eliminated from adult pancreas tissues via other routes. We also found no evidence of compensatory proliferation of normal cells and instead observed a decrease in global cell proliferation over time in all tissues, irrespective of genotype. This is consistent with previous reports demonstrating that the proliferation rate in pancreas epithelia declines with age^34,35^. Indeed, pancreas tissues are generally regarded as slow proliferating tissues. Here, we report a novel finding whereby competitive interactions in exocrine tissues lead to a change in normal and mutant cell size *in vivo*. KrasG12D cells decrease in cell volume and normal cells directly neighbouring KrasG12D cells increase in cell volume to compensate for the loss of mutant cell size. In *Drosophila melanogaster*, competition for space and apoptosis of ‘loser’ cells induce the rapid expansion of ‘winner’ cells immediately neighbouring ‘loser’ cells, independent of cell proliferation^46^. In this scenario, ‘winner’ cell expansion is governed by local changes in cell topology and a remodelling of cell-cell junctions at winner-loser cell boundaries^46^. Similarly, in post-mitotic tissues in *Drosophila melanogaster*, ‘winner’ cells compensate for loss of ‘loser’ cells via hypertrophic cell growth^47,48^. Hypertrophic ‘winner’ cells increase in cell volume and display an increase in endoreplication^48^; however, we found no evidence of endoreplication in murine pancreas tissues (not shown). Our data reveal that mutant and normal cell volume is unchanged in EphA2-depleted tissues. Future studies are required to determine the mechanisms underlying how pancreatic epithelial cells lose/gain volume *in vivo* and how EphA2 is contributing to this process.

We have previously shown that epithelial cells sense and respond to steep differences in Eph receptor expression *in vitro^19^* and *in Drosophila melanogaster in vivo*^10^ and that differential Eph signalling leads to the segregation and extrusion of RasV12 cells from normal neighbours. Here, we show that similar phenomena occur in pancreas epithelia in an EphA2-dependent manner. In pancreas, KrasG12D cells shrink in cell size and segregate from normal neighbours *in vivo* and *in vitro* prior to being eliminated and are often apically extruded from non-transformed monolayers *in vitro*. Indeed, we have previously demonstrated using MDCK cell lines that RasV12-expressing cells in direct contact with normal cells become contractile and segregate before being apically extruded^4,10^. Our data in exocrine pancreas reveal that KrasG12D cells are eliminated at the single cell level and mathematical modelling predicts that KrasG12D cells are eliminated at cell-cell boundaries where interaction with normal cells is highest. Together, our data support a model whereby EphA2 signalling at normal-mutant boundaries triggers the segregation and elimination of KrasG12D cells in adult pancreas. In addition, EphA2 signalling at normal-mutant boundaries would destabilise E-cadherin-based cell-cell adhesion by inducing cell repulsion^10,19^ and/or endocytosis of E-cadherin^49^. Indeed, imaging adherens junctions in pancreas tissues revealed that E-cadherin and p120 catenin are specifically decreased at cell-cell contacts between KRasG12D and normal cells in an EphA2-dependent manner *in vivo* and E-cadherin is predominantly intracellular in KRasG12D *in vitro*. Together, our data support a functional role of EphA2 in regulating E-cadherin localisation and stability at cell-cell contacts between KrasG12D and normal cells in pancreas epithelia.

Eph-ephrin biology in the pancreas is incompletely understood. Both A and B Ephs and ephrins are expressed in the developing pancreas^50^ and EphB signals regulate pancreas tissue branching and morphogenesis^51^ and cell fate decisions^52^. In adult tissues, EphA-ephrinA signals regulate insulin secretion from endocrine compartments^53^; however, the role of Eph-ephrin signalling in exocrine tissues is less clear. Eph-ephrins were originally identified as neural guidance molecules that play a crucial role in neurogenesis^54^. Intriguingly, other guidance molecules have been implicated in cell competition in epithelial tissues. In *Drosophila melanogaster*, Slit-Robo signals drive extrusion of polarity deficient cells by disrupting E-cadherin^55^, while semaphorin-plexin signalling promotes the elimination of damaged cells during epithelial wound repair^56^. We have previously shown in *Drosophila melanogaster* that DEph is required to eliminate RasV12 cells from developing epithelia *in vivo^10^*. Together with our current evidence demonstrating an essential role of EphA2 in adult pancreas, these observations suggest that cell-cell communication signals have important functions in maintaining epithelial tissue health and homeostasis.

Our findings imply that EphA2 is a novel tumour suppressor in pancreatic cancer. Interestingly, loss of EphA2 cooperates with *RAS* mutations to drive skin^57^ and lung^58^ tumourigenesis. Whether EphA2 is required to clear *RAS* mutant cells and preserve tissue health in other epithelial tissues remains to be investigated. Considering *KRAS* mutations drive all stages of pancreatic cancer^59^ (with codon 12 being the most prevalent, detected in >80% of human tumours^60^), our data infer that KrasG12D mutant cells must be able to override competitive signals to survive in tissues and initiate tumourigenesis. Remarkably, our study reveals that a rare population of KrasG12D cells are never eliminated and go on to initiate premalignant lesion development. An exciting line of investigation for future work will be to determine the mechanisms underlying how these rare KrasG12D cells evade competitive signals with normal cells to survive in tissues. A better understanding of the mechanisms by which normal cells outcompete KrasG12D cells could lead to therapies to alter the fitness landscape^15^, promoting elimination of aberrant cells and decreasing pancreatic cancer incidence^61^. Equally, understanding the mechanisms underlying how KrasG12D cells increase fitness could provide novel insights into how risk factors drive disease.

## Author Contributions

Conceptualization, C.H.; Methodology, W.H., A.Z., B.S., M.A., A.P., S.P., T.W.; Investigation & Analysis, W.H., A.Z., B.S., M.A., A.P., S.P., T.W., J.M., O.S., C.H.; Writing – original Draft, C.H.; Writing – Review and Editing, C.H., W.H.; Resources, G.P., T.W., J.M., O.S.; Supervision, C.H.

## Acknowledgements

We thank T. Dale (School of Biosciences, Cardiff University) for mentorship and for accommodating the mouse work on his project license. We thank F. Afonso (School of Biosciences, Cardiff University) and J. de Navascués (University of Essex) for critical reading of the manuscript and helpful discussions with data analysis. A.Z., A.P., and S.P. were supported by Amser Justin Time (registered charity 1124951). W.H. was supported by MRC DTG PhD studentship. B.S., M.A., are supported by CRUK Early detection project grant (A27838) to C.H. J.P.M and O.J.S are supported by Cancer Research UK (A25142, A17196, A21139 and A25233). C.H. is a European Cancer Stem Cell Research Institute Fellow (Cardiff University). This work was supported by Amser Justin Time funding (registered charity 1124951) and was initiated with a Pancreatic Cancer UK Research Innovation Fund (A16868) to C.H. Authors declare no conflict of interests

## Methods

### Animals and Induction of Cre recombinase *in vivo*

*Pdx1-Cre^ER^, LSL-Kras^G12D^, Rosa26^LSL-tdRFP^ and EphA2*^−/−^ mouse lines have all been previously described^1–4^. Animals were housed in conventional pathogen-free animal facilities and experiments were conducted in accordance with U.K. Home Office regulations (ASPA 1986 & EU Directive 2010) under the guidelines of Cardiff University Animal Welfare and Ethics Committee. Mice were genotyped by PCR analysis following standard methods. Details on primer sequences can be found in Supplemental Table 1. Both male and female experimental and control cohorts were injected at 6-8 weeks of age by intraperitoneal injection of tamoxifen in corn oil. Low frequency expression of RFP was induced via a single intraperitoneal injection of 1 µg/40 g bodyweight (low dose) with widespread recombination (high dose) induced by three injections of 9 mg/40 g over 5 days. At specified time points the pancreas was harvested and routinely dissected into three segments. Segments were then either fixed in 10% buffered formalin overnight at 4°C for IHC, snap frozen in OCT or fixed in 2% paraformaldehyde for immunofluorescent tomography.

### Scoring RFP in tissues

To measure global levels of endogenous RFP fluorescence, five 10 µm thick cryosections were cut from fresh frozen tissues with each slice at least 50 µm apart and immediately imaged. Whole sections were imaged on a Zeiss confocal LSM 710 using tile scans for brightfield and RFP. Global endogenous RFP fluorescence was calculated as a proportion of total tissue area. Tissue area was quantified from brightfield images and RFP fluorescence was segmented and measured using ImageJ. For cluster analysis, clusters were discretely segmented and quantified in ImageJ and binned on size before normalising to total tissue area. This produced a density of clusters of different sizes which was plotted in GraphPad. RFP positive ducts and RFP positive acinar cells were scored in tissues fixed and stained using immunofluorescence tomography protocols (described below). RFP positive islets were scored in formalin-fixed paraffin-embedded (FFPE) tissues and immunohistochemistry protocols (described below). Recombined RFP in genomic DNA (gDNA) was measured by qPCR (SyGreen mix; PCR Biosystems) and primer sets (Supplemental Table 2). gDNA extraction was carried out from tissues using the DNeasy microkit (Qiagen) following the manufacturer’s instructions. Levels of recombined RFP allele relative to ApoB were analysed by the delta Ct method.

### Immunohistochemistry (IHC) and Immunofluorescence

Histological staining was performed on FFPE sections. For RFP IHC, tissue was fixed in 10% neutral buffered formalin (Sigma-Aldrich) for 24 h at 4°C. Formalin was replaced by 70% ethanol and tissue was processed into wax blocks by standard methods. Sections were cut at 5 µm thickness, dewaxed and rehydrated. For antigen retrieval, tissue sections were incubated in 20 µg/ml Proteinase K (Roche) diluted in TBS/T for 15 min at 37°c. Sections were blocked in 3% H_2_O_2_ (Sigma Aldrich) for 20 min, followed by 5% Normal Goat Serum (NGS) (S-1000, Vector Labs) for 30 min. Anti-RFP (Rabbit, polyclonal) antibody (600-401-379, Rockland) was used at 1:500. Incubation with the primary antibody was performed overnight at 4°c. Secondary antibody ImmPRESS goat anti-Rabbit (MP-7451, Vector Labs) was added at room temperature for 30 min followed by DAB chromogen (Peroxidase substrate kit: SK-4100, Vector Labs) for 3 min. For cleaved caspase 3 and Ki67 IHC, samples were washed in xylene before rehydration in decreasing concentrations of ethanol. After washing, antigen retrieval was carried out using citrate buffer (10mM, pH 6.0) and boiling in a pressure cooker for 5 min (Ki67) or 8 min (CC3). Endogenous peroxidase was blocked by incubation in 0.5% H_2_O_2_ for 20 min (Ki67) or 3% H_2_O_2_ for 10 min (CC3). Sections were blocked in 20% NGS for 30 min (Ki67) or 5% NGS for 60 min (CC3) before incubation in primary antibody overnight (anti-Ki67, Abcam; anti-cleaved Caspase 3, Cell Signalling). Following washing, samples were incubated with biotinylated secondaries (E0432, DAKO) before visualisation using Vectastain ABC kit (PK-6100, Vector Lab) and counterstained with DAB. IHC stained tissues were scanned using the Axio Scan Z1 slide scanner (Zeiss, Cambridge, UK), using a 20X magnification, and images were analysed using the Zeiss Axio Scan Zen software. Quantification of IHC (cells positive for antibody staining, or proportion of RFP positive islet cells) was carried out manually using Zeiss Zen software. For Alcian Blue staining, FFPE tissue sections were dewaxed in xylene and rehydrated in decreasing concentrations of ethanol, before washing in 3% acetic acid for 5 min. Tissue sections were incubated in staining solution (1% Alcian Blue in 3% acetic acid pH 2.5) for 10 min, before extensively washed in 3% acetic acid. Following water rinsing, tissue sections were counterstained using Nuclear Fast Red solution for 10 min (N3020, Sigma, Nuclear Fast Red: 0.1% in 5% aluminium sulphate). Following staining, specimens were rinsed in water, dehydrated and equilibrated into xylene, and mounted with DPX mountant. The entire tissue area was imaged on a Zeiss Axio Scan Z1 slide scanner followed by surface area measurement using Zeiss ZEN software. Alcian Blue positive PanIN lesions per mm^2^ surface area were counted manually. TUNEL staining was carried out on FFPE tissues using a commercially available kit (Abcam) according to the manufacturer’s protocol and Hoescht 33342. Stained tissues were imaged using a Leica DMI600B inverted epifluorescence microscope. Primary antibodies are detailed in Supplemental Table 3.

### Immunofluorescence tomography (IT)

Immunofluorescence tomography (IT) is a high-resolution 3D reconstruction method based on embedding in methacrylate followed by serial sectioning^5^. Computational alignment of 2D immunostained serial sections produces a 3D volume rendering and was carried out as previously described^5,6^. Briefly, fixed tissue was embedded in butyl-methyl methacrylate plastic (BMMA) under UV following dehydration and resin infiltration. Serial 2 pm-thick sections were then rehydrated, and antigens unmasked by boiling in citrate buffer for at least 7 min. Samples were blocked in 5% NGS before incubation in primary antibody overnight at 4°c. After three PBS washes, samples were incubated for two hours at room temperature in appropriate secondary antibodies (1:200; Life Technologies) before washing, staining nuclei with Hoescht 33342 and mounting in Mowiol. Individual sections were then imaged and aligned semi-automatically using Amira (Version 5.4).

### Primary pancreatic ductal epithelial cells (PDEC) and coculture assays

Isolation of transformed mouse pancreatic cells was previously described^7^. Transformed cells were maintained in Dulbecco’s modified Eagle medium (DMEM) supplemented with 10% FBS and 1% penicillin/streptomycin. Normal primary cells were established as described previously^8^. Briefly, whole pancreas was mechanically dissociated before digestion in collagenase at 37°c. Following several washes in HBSS supplemented with 5% FBS, tissue was passed through a 40 µm cell strainer. Cells were then digested with trypsin for 5 min before washing to remove any trace of collagenase. The washed cells were then resuspended in PDEC media (DMEM/F12 with 5 % Nu-Serum, 5 mg/ml glucose, 1.22 mg/ml nicotinamide, 0.1 mg/ml soybean trypsin inhibitor, 25 mg/ml bovine pituitary extract, 20 ng/ml EGF, 100 ng/ml cholera toxin, 5 nM 3,3,5-tri-iodo-L-thyronine, 5 mL/L ITS+ culture supplement, 1 mM dexamethasone). Cells were plated on rat collagen I-coated 6-well plates. Cell-cell mixing experiments and staining of cells was carried out as described previously^9,10^. In brief, one cell population was labelled with CMRA cell tracker dye (ThermoFisher Scientific) as previously described^9^, before mixing with unlabelled cells at 1:50 ratios. After 48 hours, cells were fixed in 4% PFA for 15 min, before 15 min permeabilization in 0.25% Triton X-100/PBS and blocking in 3% BSA/PBS for 1 hour. Cells were then incubated in primary antibody (detailed in Supplemental Table 3), diluted in blocking buffer, overnight at 4°C. The following day, cells were washed three times with PBS and incubated in secondary antibodies (1:200, Life Technologies) and phalloidin (1:200, Invitrogen) for 1 h at room temperature. Cells were then mounted in Mowiol and imaged on a Zeiss confocal LSM 710.

### Image analysis

Inter-nuclear distance (IND) was calculated by measuring the distance between the centre of nuclei of neighbouring cells in 3D projections using Imaris (Bitplane, Version 8.0). Nuclei were initially segmented using ‘spots’ before measuring from the centre of each spot. To measure cluster area and cell volume, cells were first segmented using Amira. To measure cluster area and index of sphericity, clusters were manually segmented using Fiji (Version 2.0) and roundness was calculated as previously described^10^. Quantification of fluorescent signal for E-cadherin and p120 catenin regions of interest was carried out in ImageJ. Images were taken under the same exposure time and settings and the mean signal intensity was measured.

### Statistics

For data analyses, normally distributed data, as determined by the Shapiro-Wilke test was analysed using unpaired Student t-tests with Welch correction. Data that were not normally distributed were compared using non-parametric Mann Whitney U two-tailed t tests. A *p* value of <0.05 was taken as significant. No statistical method was used to pre-determine sample size. For animal studies, experiments were not randomised, and investigators were not blinded to allocation during experiments. All experiments were reproduced with at least three independent experiments and at least three animals of each genotype.

### Mathematical Model

We consider two spatial populations denoted by ***m*** for mutant and ***w*** for wild type. These two populations interact on a square based grid, so each point is inhabited by at most one tissue type and the position can be specified by a discrete lattice coordinate system. Namely, each lattice site is a square of length ***δ*** and the ***(i j)^th^*** lattice site is either occupied by a wild type tissue (i.e.,***m(i, j)*** = **0**,***w(i, j)*** = **1**) or a mutant tissue type (i.e.,***m(i, j)*** = **1**,***w(i, j)*** = **0**). Whenever a mutant cell is next to a wild type cell the two cell types compete, with the wild type cell winning at a rate d. Critically, we only consider two cells as neighbours if they share a boundary. Thus, the cells only interact with their north, east, south and west neighbours (see SFig. 4A).

By considering a general spatial site (*i, J*) we write down all possible combinations of actions that can occur at that site. Specifically,

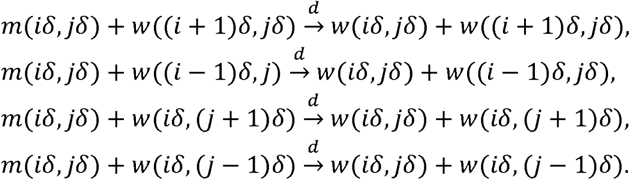

The original interaction equations are simulated using a standard "Gillespie” Stochastic Simulation Algorithm (SSA)^11–13^. To use a SSA we first need to calculate the propensity, *a*_*r*_, of each reaction, *r*, where the propensity is the probability per unit time that a specific reaction occurs. Further, we calculate the probability per unit time that any reaction occurs, which is the sum of all propensities,

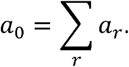

The SSA proceeds as follows:

1. Define the initial time, *t* = 0, and the final time, *t*_*f*_, over which we wish to run the algorithm.
2. Generate two uniformly randomly distributed numbers, *r*_*1*_ and *r*_*2*_, from the interval [0, 1].
3. Calculate all propensity functions, *a*_*r*_, and their sum *a*_*0*_.
4. Compute the time, τ, when the next reaction takes place, where

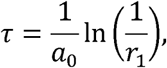

and update the current time, *t*: = *t* + τ.
5. Find the reaction that fires by searching for the integer, *j*, such that

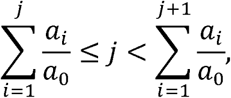

and update the populations based on the reaction that fires.
6. If *t* ≥ *t*_*f*_ then end the simulation, otherwise go to step 2.

This algorithm allows us to accurately simulate a single stochastic trajectory with the correct probabilistic properties. Simulating multiple trajectories allows us to produce probability distributions that provide us with the probability of the system inhabiting a given state at a given time.

---

**Supplemental Table 1:**
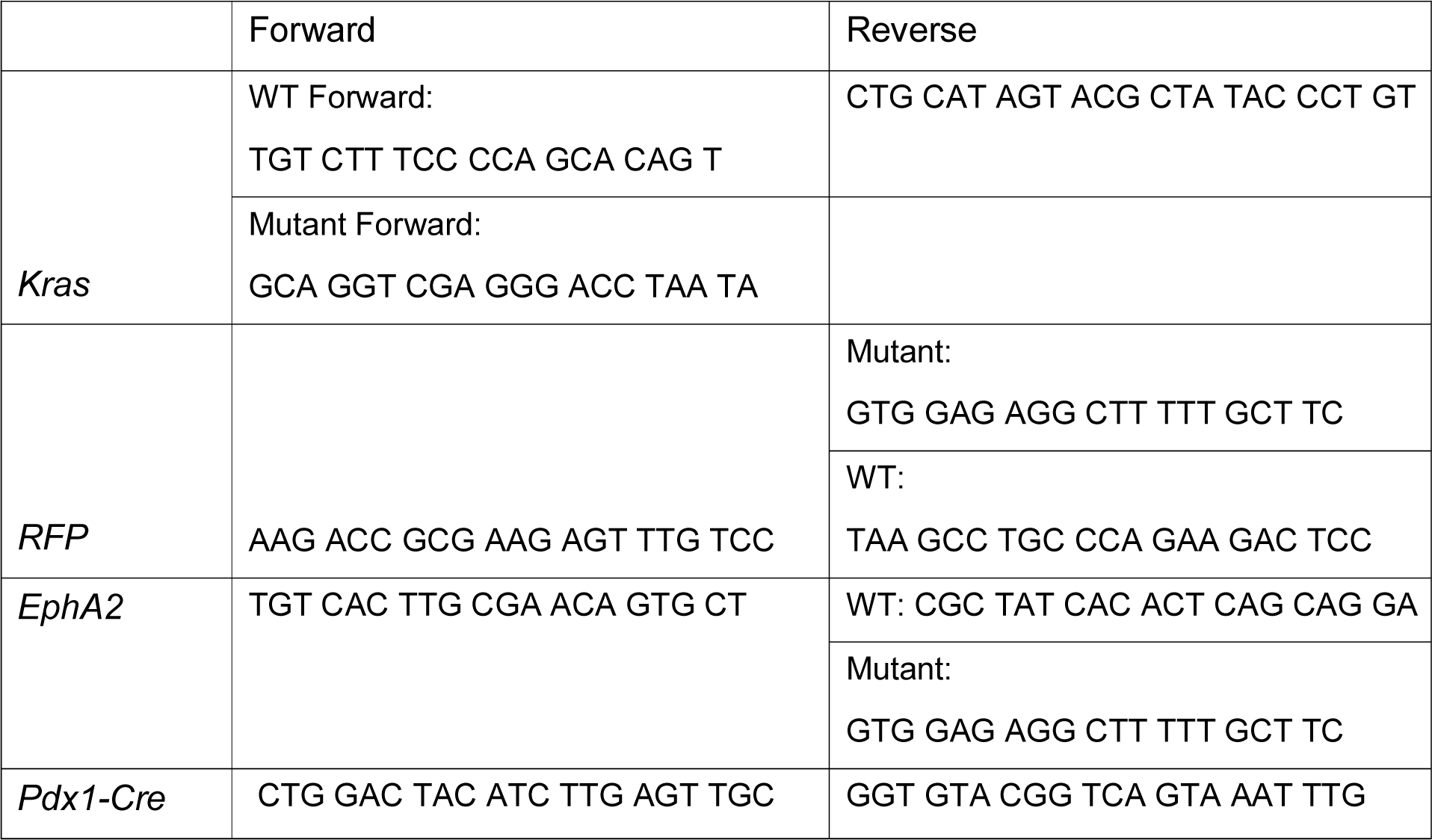
Genotyping Primers.

**Supplemental Table 2:**
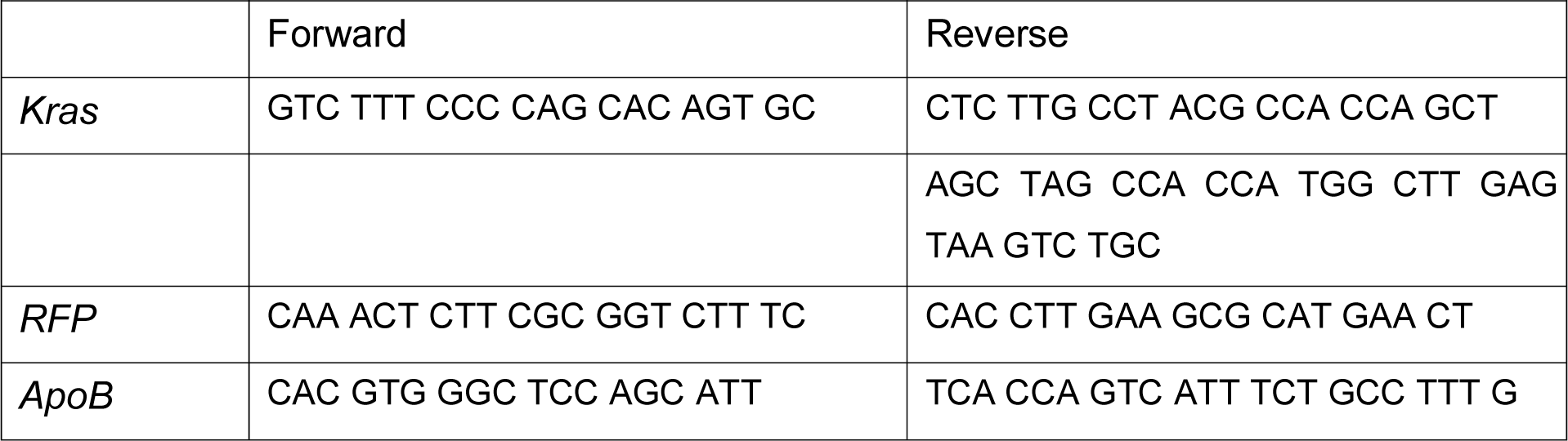
Recombined alleles (gDNA)

**Supplemental Table 3:**
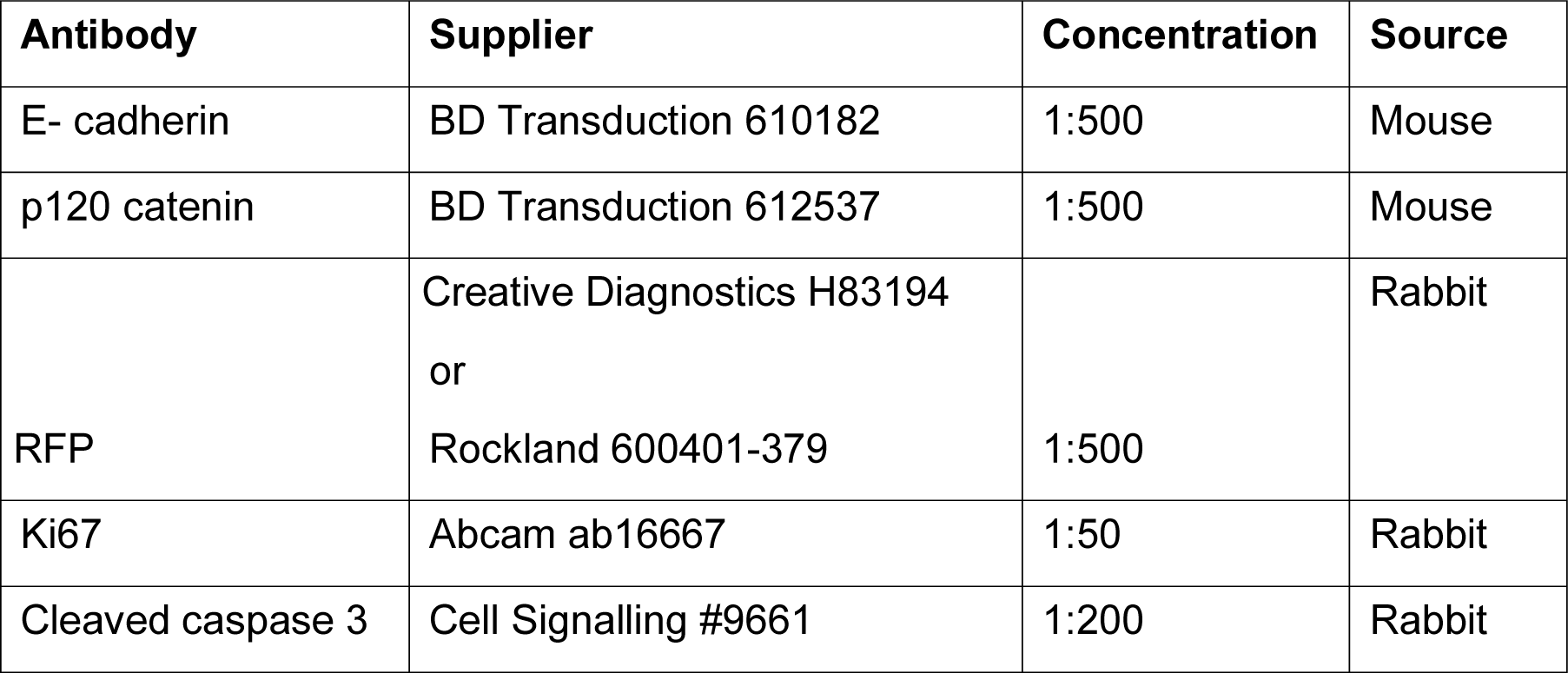
Primary antibodies.

## Notes

### Competing Interest Statement

The authors have declared no competing interest.

## References

1. Eisenhoffer, G. T. et al. Crowding induces live cell extrusion to maintain homeostatic cell numbers in epithelia. Nature 484, 546–549, doi:10.1038/naturel0999 (2012).

2 Rosenblatt, J., Raff, M. C. & Cramer, L. P. An epithelial cell destined for apoptosis signals its neighbors to extrude it by an actin- and myosin-dependent mechanism. Curr Biol 11, 1847–1857, doi:10.1016/s0960-9822(01)00587-5 (2001).

3 Brown, S. et al. Correction of aberrant growth preserves tissue homeostasis. Nature 548, 1334–337, doi:10.1038/nature23304 (2017).

4 Hogan, C. et al. Characterization of the interface between normal and transformed epithelial cells. Nat Cell Biol 11, 460–467, doi:10.1038/ncbl853 (2009).

5 Kajita, M. et al. Interaction with surrounding normal epithelial cells influences signalling pathways and behaviour of Src-transformed cells. J Cell Sci 123, 171–180, doi: 10.1242/jcs.057976 (2010).

6 Kon, S. et al. Cell competition with normal epithelial cells promotes apical extrusion of transformed cells through metabolic changes. Nat Cell Biol 19, 530–541, doi:10.1038/ncb3509 (2017).

7 Murai, K. et al. Epidermal Tissue Adapts to Restrain Progenitors Carrying Clonal p53 Mutations. Cell Stem Cell 23, 687–699 e688, doi:10.1016/j.stem.2018.08.017 (2018).

8 Norman, M. et al. Loss of Scribble causes cell competition in mammalian cells. J Cell Sci 125, 59–66, doi: 10.1242/jcs.085803 (2012).

9 Pineda, C. M. et al. Hair follicle regeneration suppresses Ras-driven oncogenic growth. J Cell Biol 218, 3212–3222, doi:10.1083/jcb.201907178 (2019).

10 Porazinski, S. et al. EphA2 Drives the Segregation of Ras-Transformed Epithelial Cells from Normal Neighbors. Curr Biol 26, 3220–3229, doi:10.1016/j.cub.2016.09.037 (2016).

11 Sasaki, A. et al. Obesity Suppresses Cell-Competition-Mediated Apical Elimination of RasV12-Transformed Cells from Epithelial Tissues. Cell Rep 23, 974–982, doi: 10.1016/j.celrep.2018.03.104 (2018).

12 Sato, N. et al. The COX-2/PGE2 pathway suppresses apical elimination of RasV12- transformed cells from epithelia. Commun Biol 3, 132, doi:10.1038/s42003-020-0847-y (2020).

13 Gu, Y. et al. Defective apical extrusion signaling contributes to aggressive tumor hallmarks. Elife 4, e04069, doi:10.7554/eUfe.04069 (2015).

14 Di Gregorio, A., Bowling, S. & Rodriguez, T. A. Cell Competition and Its Role in the Regulation of Cell Fitness from Development to Cancer. Dev Cell 38, 621–634, doi: 10.1016/j.devcel.2016.08.012 (2016).

15 Vishwakarma, M. & Piddini, E. Outcompeting cancer. Nat Rev Cancer 20, 187–198, doi:10.1038/s41568-019-0231-8 (2020).

16 Matamoro-Vidal, A. & Levayer, R. Multiple Influences of Mechanical Forces on Cell Competition. Curr Biol 29, R762–R774, doi:10.1016/j.cub.2019.06.030 (2019).

17 Wagstaff, L. et al. Mechanical cell competition kills cells via induction of lethal p53 levels. Nat Commun 7, 11373, doi:10.1038/ncommsll373 (2016).

18 Tai, K., Cockburn, K. & Greco, V. Flexibility sustains epithelial tissue homeostasis. Curr Opin Cell Biol 60, 84–91, doi:10.1016/j.ceb.2019.04.009 (2019).

19 Hill, W. & Hogan, C. Normal epithelial cells trigger EphA2-dependent RasV12 cell repulsion at the single cell level. Small GTPases 10, 305–310, doi:10.1080/21541248.2017.1324940 (2019).

20 Lisabeth, E. M., Falivelli, G. & Pasquale, E. B. Eph receptor signaling and ephrins. Cold Spring Harb Perspect Biol 5, doi:10.1101/cshperspect.a009159 (2013).

21 Batlle, E. & Wilkinson, D. G. Molecular mechanisms of cell segregation and boundary formation in development and tumorigenesis. Cold Spring Harb Perspect Biol 4, a008227, doi:10.1101/cshperspect.a008227 (2012).

22 Demcollari, T. I., Cujba, A. M. & Sancho, R. Phenotypic plasticity in the pancreas: new triggers, new players. Curr Opin Cell Biol 49, 38–46, doi:10.1016/j.ceb.2017.11.014 (2017).

23 Buscail, L., Bournet, B. & Cordelier, P. Role of oncogenic KRAS in the diagnosis, prognosis and treatment of pancreatic cancer. Nat Rev Gastroenterol Hepatol 17, 153–168, doi:10.1038/s41575-019-0245-4 (2020).

24 Chari, S. T. et al. Early detection of sporadic pancreatic cancer: summative review. Pancreas 44, 693–712, doi:10.1097/MPA.0000000000000368 (2015).

25 Guerra, C. et al. Chronic pancreatitis is essential for induction of pancreatic ductal adenocarcinoma by K-Ras oncogenes in adult mice. Cancer Cell 11, 291–302, doi:10.1016/j.ccr.2007.01.012 (2007).

26 Morton, J. P. et al. Mutant p53 drives metastasis and overcomes growth arrest/senescence in pancreatic cancer. Proc Natl Acad Sci USA 107, 246–251, doi:10.1073/pnas.0908428107 (2010).

27 Cooper, C. L., O'Toole, S. A. & Kench, J. G. Classification, morphology and molecular pathology of premalignant lesions of the pancreas. Pathology 45, 286–304, doi:10.1097/PAT.0b013e32835f2205 (2013).

28 Gidekel Friedlander, S. Y. et al. Context-dependent transformation of adult pancreatic cells by oncogenic K-Ras. Cancer Cell 16, 379–389, doi:10.1016/j.ccr.2009.09.027 (2009).

29 Ferreira, R. M. M. et al. Duct- and Acinar-Derived Pancreatic Ductal Adenocarcinomas Show Distinct Tumor Progression and Marker Expression. Cell Rep 21, 966–978, doi:10.1016/j.celrep.2017.09.093 (2017).

30 Kopp, J. L. et al. Identification of Sox9-dependent acinar-to-ductal reprogramming as the principal mechanism for initiation of pancreatic ductal adenocarcinoma. Cancer Cell 22, 737–750, doi:10.1016/j.ccr.2012.10.025 (2012).

31 Kopp, J. L., Grompe, M. & Sander, M. Stem cells versus plasticity in liver and pancreas regeneration. Nat Cell Biol 18, 238–245, doi:10.1038/ncb3309 (2016).

32 Varga, J. & Greten, F. R. Cell plasticity in epithelial homeostasis and tumorigenesis. Nat Cell Biol 19, 1133–1141, doi:10.1038/ncb3611 (2017).

33 Hingorani, S. R. et al. Preinvasive and invasive ductal pancreatic cancer and its early detection in the mouse. Cancer Cell 4, 437–450, doi:10.1016/sl535-6108(03)00309-x (2003).

34 Houbracken, I. & Bouwens, L. Acinar cells in the neonatal pancreas grow by self-duplication and not by neogenesis from duct cells. Sci Rep 7, 12643, doi:10.1038/s41598-017-12721-9 (2017).

35 Teta, M., Rankin, M. M., Long, S. Y., Stein, G. M. & Kushner, J. A. Growth and regeneration of adult beta cells does not involve specialized progenitors. Dev Cell 12, 817–826, doi: 10.1016/j.devcel.2007.04.011 (2007).

36 Brantley-Sieders, D. M. et al. EphA2 receptor tyrosine kinase regulates endothelial cell migration and vascular assembly through phosphoinositide 3-kinase-mediated Rad GTPase activation. J Cell Sci 117, 2037–2049, doi:10.1242/jcs.01061 (2004).

37 Parfitt, G. J. Immunofluorescence Tomography: High-resolution 3-D reconstruction by serial-sectioning of methacrylate embedded tissues and alignment of 2-D immunofluorescence images. Sci Rep 9, 1992, doi:10.1038/s41598-018-38232-9 (2019).

38 Parfitt, G. J. et al. A novel immunofluorescent computed tomography (ICT) method to localise and quantify multiple antigens in large tissue volumes at high resolution. PLoS One 7, e53245, doi:10.1371/journal.pone.0053245 (2012).

39 Nanes, B. A. et al. pl20-catenin binding masks an endocytic signal conserved in classical cadherins. J Cell Biol 199, 365–380, doi:10.1083/jcb.201205029 (2012).

40 Bulgakova, N. A. & Brown, N. H. Drosophila pl20-catenin is crucial for endocytosis of the dynamic E-cadherin-Bazooka complex. J Cell Sci 129, 477–482, doi:10.1242/jcs.177527 (2016).

41 Bowling, S., Lawlor, K. & Rodriguez, T. A. Cell competition: the winners and losers of fitness selection. Development 146, doi: 10.1242/dev.167486 (2019).

42 Kräh, N. M. et al. The acinar differentiation determinant PTF1A inhibits initiation of pancreatic ductal adenocarcinoma. Elife 4, doi:10.7554/eLife.07125 (2015).

43 Merino, M. M. et al. Elimination of unfit cells maintains tissue health and prolongs lifespan. Cell 160, 461–476, doi:10.1016/j.cell.2014.12.017 (2015).

44 Martins, V. C. et al. Cell competition is a tumour suppressor mechanism in the thymus. Nature 509, 465–470, doi:10.1038/naturel3317 (2014).

45 Ellis, S. J. et al. Distinct modes of cell competition shape mammalian tissue morphogenesis. Nature 569, 497–502, doi:10.1038/s41586-019-1199-y (2019).

46 Tsuboi, A. et al. Competition for Space Is Controlled by Apoptosis-Induced Change of Local Epithelial Topology. Curr Biol 28, 2115–2128 e2115, doi:10.1016/j.cub.2018.05.029 (2018).

47 Tamori, Y. & Deng, W. M. Compensatory cellular hypertrophy: the other strategy for tissue homeostasis. Trends Cell Biol 24, 230–237, doi:10.1016/j.tcb.2013.10.005 (2014).

48 Tamori, Y. & Deng, W. M. Tissue repair through cell competition and compensatory cellular hypertrophy in postmitotic epithelia. Dev Cell 25, 350–363, doi:10.1016/j.devcel.2013.04.013 (2013).

49 Saitoh, S. et al. Rab5-regulated endocytosis plays a crucial role in apical extrusion of transformed cells. Proc Natl Acad Sci USA 114, E2327–E2336, doi:10.1073/pnas.1602349114 (2017).

50 van Eyll, J. M. et al. Eph receptors and their ephrin ligands are expressed in developing mouse pancreas. Gene Expr Patterns 6, 353–359, doi:10.1016/j.modgep.2005.09.010 (2006).

51 Villasenor, A., Chong, D. C., Henkemeyer, M. & Cleaver, O. Epithelial dynamics of pancreatic branching morphogenesis. Development 137, 4295–4305, doi:10.1242/dev.052993 (2010).

52 Villasenor, A. et al. EphB3 marks delaminating endocrine progenitor cells in the developing pancreas. Dev Dyn 241, 1008–1019, doi:10.1002/dvdy.23781 (2012).

53 Konstantinova, I. et al. EphA-Ephrin-A-mediated beta cell communication regulates insulin secretion from pancreatic islets. Cell 129, 359–370, doi:10.1016/j.cell.2007.02.044 (2007).

54 Kania, A. & Klein, R. Mechanisms of ephrin-Eph signalling in development, physiology and disease. Nat Rev Mol Cell Biol 17, 240–256, doi:10.1038/nrm.2015.16 (2016).

55 Vaughen, J. & Igaki, T. Slit-Robo Repulsive Signaling Extrudes Tumorigenic Cells from Epithelia. Dev Cell 39, 683–695, doi:10.1016/j.devcel.2016.11.015 (2016).

56 Yoo, S. K. et al. Plexins function in epithelial repair in both Drosophila and zebrafish. Nat Commun 7, 12282, doi:10.1038/ncommsl2282 (2016).

57 Guo, H. et al. Disruption of EphA2 receptor tyrosine kinase leads to increased susceptibility to carcinogenesis in mouse skin. Cancer Res 66, 7050–7058, doi:10.1158/0008-5472.CAN-06-0004 (2006).

58 Yeddula, N., Xia, Y., Ke, E., Beumer, J. & Verma, I. M. Screening for tumor suppressors: Loss of ephrin receptor A2 cooperates with oncogenic KRas in promoting lung adenocarcinoma. Proc Natl Acad Sci U S A 112, E6476–6485, doi:10.1073/pnas.l520110112 (2015).

59 Collins, M. A. et al. Oncogenic Kras is required for both the initiation and maintenance of pancreatic cancer in mice. J Clin Invest 122, 639–653, doi:10.1172/JCI59227 (2012).

60 Lanfredini, S., Thapa, A. & O'Neill, E. RAS in pancreatic cancer. Blochern Soc Trans 47, 961–972, doi:10.1042/BST20170521 (2019).

61 Fernandez-Antoran, D. et al. Outcompeting p53-Mutant Cells in the Normal Esophagus by Redox Manipulation. Cell Stem Cell 25, 329–341 e326, doi:10.1016/j.stem.2019.06.011 (2019).

## Supplemental bibliography

1 Brantley-Sieders, D. M. et al. EphA2 receptor tyrosine kinase regulates endothelial cell migration and vascular assembly through phosphoinositide 3-kinase-mediated GTPase activation. J Cell Sci 117, 2037–2049, doi:10.1242/jcs.01061 (2004).

2 Gu, G., Dubauskaite, J. & Melton, D. A. Direct evidence for the pancreatic lineage: NGN3+ cells are islet progenitors and are distinct from duct progenitors. Development 129, 2447–2457 (2002).

3 Hingorani, S. R. et al. Preinvasive and invasive ductal pancreatic cancer and its early detection in the mouse. Cancer Cell 4, 437–450, doi:10.1016/s1535-6108(03)00309-x (2003).

4 Luche, H., Weber, O., Nageswara Rao, T., Blum, C. & Fehling, H. J. Faithful activation of an extra-bright red fluorescent protein in “knock-in” Cre-reporter mice ideally suited for lineage tracing studies. Eur J Immunol 37, 43–53, doi:10.1002/eji.200636745 (2007).

5 Parfitt, G. J. et al. A novel immunofluorescent computed tomography (ICT) method to localise and quantify multiple antigens in large tissue volumes at high resolution. PLoS One 7, e53245, doi:10.1371/journal.pone.0053245 (2012).

6 Parfitt, G. J. Immunofluorescence Tomography: High-resolution 3-D reconstruction by serial-sectioning of methacrylate embedded tissues and alignment of 2-D immunofluorescence images. Sci Rep 9, 1992, doi:10.1038/s41598-018-38232-9 (2019).

7 Morton, J. P. et al. Mutant p53 drives metastasis and overcomes growth arrest/senescence in pancreatic cancer. Proc Natl Acad Sci USA 107, 246–251, doi:10.1073/pnas.0908428107 (2010).

8 Means, A. L. et al. Pancreatic epithelial plasticity mediated by acinar cell transdifferentiation and generation of nestin-positive intermediates. Development 132, 3767–3776, doi: 10.1242/dev.01925 (2005).

9 Hogan, C. et al. Characterization of the interface between normal and transformed epithelial cells. Nat Cell Biol 11, 460–467, doi:10.1038/ncb1853 (2009).

11 Gillespie, D. T. Exact stochastic simulation of coupled chemical reactions. J. Phys. Chem. 81, 2340–2361, doi:https://doi.org/10.1021/ilQ0540a008 (1977).

12 Gillespie, D. T. A general method for numerically simulating stochastic time evolution of coupled chemical reactions. J. Comput. Phys. 22, 403–434 (1976).

13 Gillespie, D., T.Concerning the validity of the stochastic approach to chemical kinetics. J. Stat. Phys. 16, 311–318, doi:https://doi.org/10.1007/BF01020385 (1977).

