## Supplemental legends for "KRASG12D mutant cells are outcompeted by wild type neighbours in adult pancreas in an EPHA2-dependent manner"

**SFigure 1: Clearance of KrasG12D cells occurs throughout the pancreas and requires the presence of normal cells.** (**A**) Bar graph depicting proportion of endogenous RFP fluorescence per tissue area in *Kras* wild type control tissues harvested at 7 days p.i following no tamoxifen (uninduced; orange bar), low dose (yellow bar) or high dose (beige bar) tamoxifen. Data represent mean +/- s.d. from 2 mice/treatment. (**B**) Scatter plot showing percentage endogenous RFP fluorescence per tissue area in *Kras* wild type control tissues harvested at 7 days p.i. Data represent mean +/- s.d. of fluorescence averaged from two mice. Tissue sections were sampled every 250 μm from the head to the tail of the pancreas. (**C**) Representative stitched confocal tile scan images of fresh frozen murine pancreas tissues showing endogenous RFP fluorescence. Tissues were harvested from control (*Kras* wild type; top panels) or KrasG12D (lower panels) at 7 days (left panels) or 35 days (right panels) p.i. following high dose of tamoxifen. (**D**) Bar graph showing percentage endogenous RFP fluorescence per tissue area in control (*Kras* wild type; white) or KrasG12D (black) tissues, harvested at 7 days (circles) or 35 days (squares) p.i. following high dose of tamoxifen. Data represent mean +/- s.d. of five tissue slices (of 50 μm apart) per mouse. n.s.=not significant. n=3 mice for control (7 days, 35 days); n=3 mice KrasG12D (7 days) and n=4 mice KrasG12D (35 days). Scale bar, 500 μm.

**SFigure 2: Cell death and cell proliferation events in adult pancreas tissues following low dose tamoxifen induction.** (**A**) Representative epifluorescence images of TUNEL assays. Positive control: mouse mammary tumour tissue; Kras WT: *Kras* wild type control. Kras WT and KrasG12D tissues were fixed at 7 days p.i. following low dose tamoxifen. Scale bar, 100 μm. (**B**-**E**) Scatter plots showing number of cells positive for (**B**) cleaved caspase 3 or (**C**) Ki67 positive cells per tissue area in control (Kras wild type; white) and KrasG12D (black) harvested at 7 days (circles) and 35 days (squares) p.i. Data represent mean +/- s.d. n.s.=not significant; **p<0.003, unpaired Student t tests with Welch correction. For (B), Control: n=7 (7 days) and n=5 (35 days) mice; KrasG12D: n=6 (7 days) and n=4 (35 days) mice. For (C), Control: n=4 (7 days) and n=5 (35 days) mice; KrasG12D: n=7 (7 days) and n=4 (35 days) mice. Scatter plots showing number of cells positive for (**D**) cleaved caspase 3 or (**E**) Ki67 positive cells per tissue area in *EphA2*^-/-^ control (*Kras* WT) (light blue) and KrasG12D EphA2-/- tissues (blue) harvested at 7 days (circles) and 35 days (squares) p.i. Data represent mean +/- s.d. n.s.=not significant; *p<0.05, **p=0.0035, unpaired Student t tests with Welch correction. For (D), EphA2-/- control: n=5 (7 days) and n=7 (35 days) mice; KrasG12D EphA2-/-: n=5 (7 days) and n=6 (35 days) mice. For (E), EphA2-/- control: n=5 (7 days) and n=7 (35 days) mice; KrasG12D EphA2-/-: n=5 (7 days) and n=6 (35 days) mice.

**SFigure 3: EphA2 expressed on KrasG12D cells drives cell segregation and apical extrusion of mutant cells from normal epithelial monolayers *in vitro*.** (**A**) Confocal images of pancreatic ductal epithelial cell (PDEC) coculture assays. Transformed tumour-derived epithelial cells (KR: KrasG12D; KRE: KrasG12D, EphA2-/-) prelabelled with cell tracker dye (CMRA: red) and mixed with non-labelled, non-transformed PDECs (N) (KR:N, KRE:N) or non-labelled transformed cells (KR:KR, KRE:KRE) at 1:50 ratios. Cells were fixed 48 h later and stained with phalloidin to visualise F-actin (grey), and Hoescht to visualise nuclei (blue). (**B**, **C**) Scatter plots of (**B**) transformed cell cluster area (μm^2^) in coculture assays, (**C**) index of sphericity of boundaries between clusters of labelled transformed cells and non-labelled neighbours. Red lines denote median. Data represent counts from n=3 repeats. ****p<0.0001, non-parametric Student t test. In (B), KR:KR: n=23; KR:N: n=36; KRE:N: n=23; KRE:KRE: n=24 clusters. In (C), KR:KR: n=23; KR:N: n=36; KRE:N: n=24; KRE:KRE: n=22 clusters (**D**) Representative confocal images of coculture assays showing KrasG12D cells are apically extruded from non-transformed cell monolayers in an EphA2-dependent manner. Transformed tumour-derived epithelial cells (KR: KrasG12D; KRE: KrasG12D, EphA2-/-) prelabelled with cell tracker dye (CMRA: red) and mixed with non-labelled, non-transformed PDECs (N) (KR:N, KRE:N) or non-labelled transformed cells (KR:KR, KRE:KRE) at 1:50 ratios. Cells were fixed 48 h later and stained with phalloidin to visualise F-actin (grey), and Hoescht to visualise nuclei (blue). White arrows label apically extruded cells. (**E**) Representative confocal images of coculture assays. Transformed tumour-derived epithelial cells (KR: KrasG12D; KRE: KrasG12D, EphA2-/-) prelabelled with cell tracker dye (CMRA: red) and mixed with non-labelled, non-transformed PDECs (KR:N, KRE:N) or non-labelled transformed cells (KR:KR, KRE:KRE) at 1:50 ratios. Cells were fixed 48 h later and stained with anti-E-cadherin antibodies (cyan) and Hoescht (blue). Scale bars, 20 μm.

**SFigure 4: Small clusters of RFP positive cells are less frequent in KC tissues over time in an EphA2-dependent manner.** (**A**) Schematic of mathematical model illustrating dynamics of normal-mutant cell interactions with normal cells in yellow and mutant cells in blue. Top panels: mutant cell with four neighbours (middle) are eliminated more readily than mutant cells with fewer normal neighbours (top right). Lower panels: simulation of the model (100x100 grid; mutant cells in blue occupy circle with radius of 50) over three time points. (**B**) Simulations of 100x100 grid as in (A, lower panels) with the radius of mutant cells altered. The initial proportion of wild-type tissue increases as the proportion of mutant tissue decreases. (**C**)-(**F**) Frequency distribution graphs of RFP positive clusters of varying size (mm^-2^). (**C**) Cluster density in *Kras* wild-type control tissues does not vary between 7 days (yellow curve) and 35 days (light grey curve). (**D**) Density of small clusters (<2000μm^2^) decreases in KrasG12D tissues at 35 days (black curve) compared to 7 days (grey curve). Cluster density in (**E**) EphA2-/-control tissues and in (**F**) KRasG12D EphA2-/- tissues does not vary between 7 days and 35 days. Data are mean +/- s.d. of minimum of 12000 clusters per genotype pooled from n=5 mice (7 days and 35 days) *Kras* wild type controls; n=4 mice (7 days) and n=6 mice (35 days) for KrasG12D; n=5 mice (7 days) and n=6 mice (35 days) for EphA2-/- controls; n=4 mice (7 days) and n=5 mice (35 days) for KrasG12D EphA2-/-. (**G**) Table of p values comparing clusters of different sizes between *Kras* wild-type (WT) and KrasG12D (KC), or KC and KrasG21D, EphA2-/- (KCE) tissues. Data were compared using unpaired Student t test with Welch correction or non-parametric Student t test, depending on results of normality tests. p<0.05 was taken as significant. Values labelled in blue indicate significance.

**Supplementary Movie legend**

**SMovie 1: Visualising RFP positive acinar cells in 3D tissues.** Representative movie of stacked confocal images of murine pancreas tissues. Using immunofluorescence tomography (IT) protocols, fixed tissues were cut into serial sections (ribbons), immunostained with anti-E-cadherin (green) and anti-RFP (red) antibodies, and Hoescht (blue) and imaged by confocal microscopy. Images were aligned and stacked to create 3D tissues using Imaris software. Movie depicts KrasG12D (KC) tissue fixed at 7 days post tamoxifen induction.
