## Supplemental Fig 1-4 for "KRASG12D mutant cells are outcompeted by wild type neighbours in adult pancreas in an EPHA2-dependent manner"

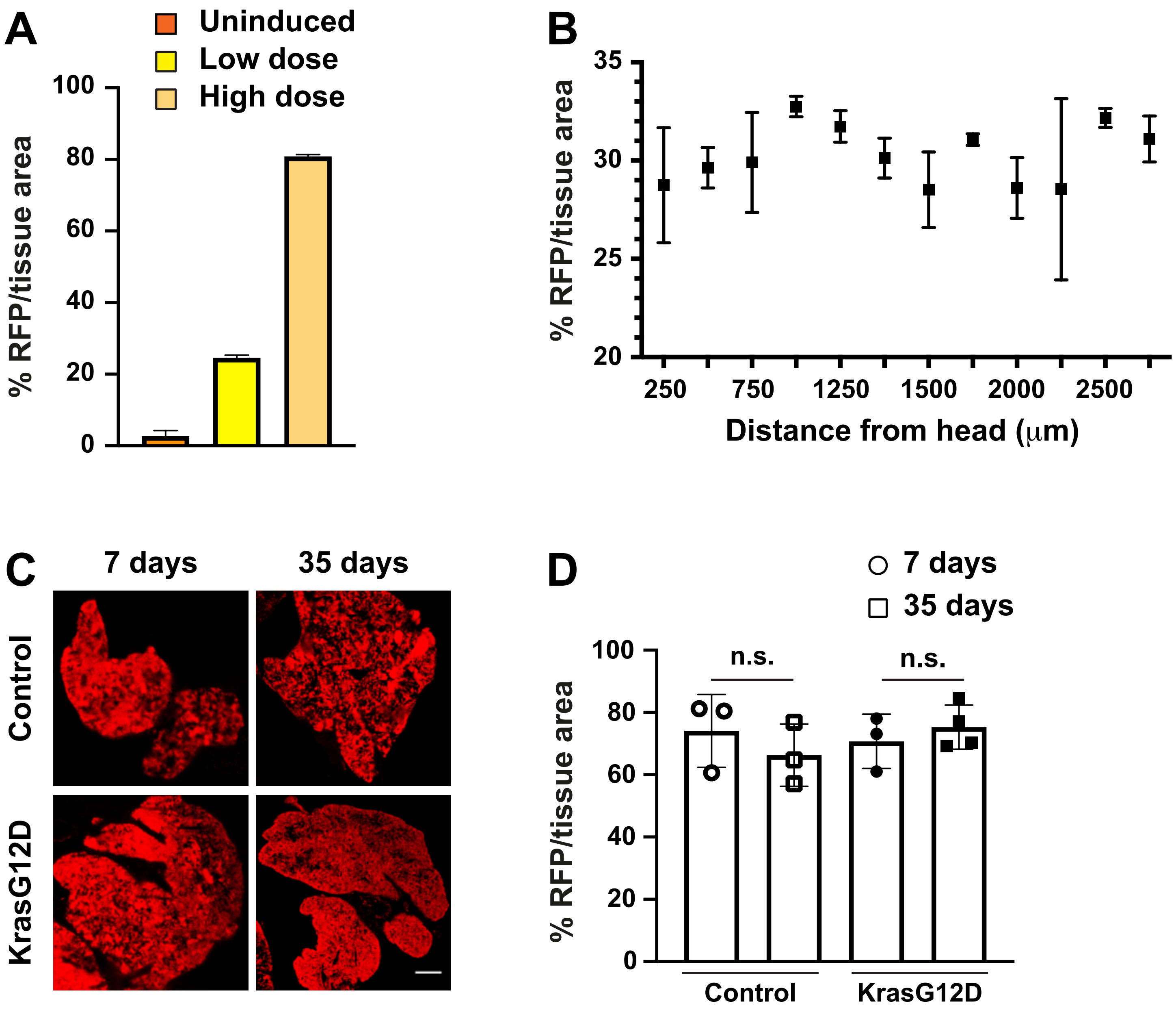


Hill et al
SFig 1

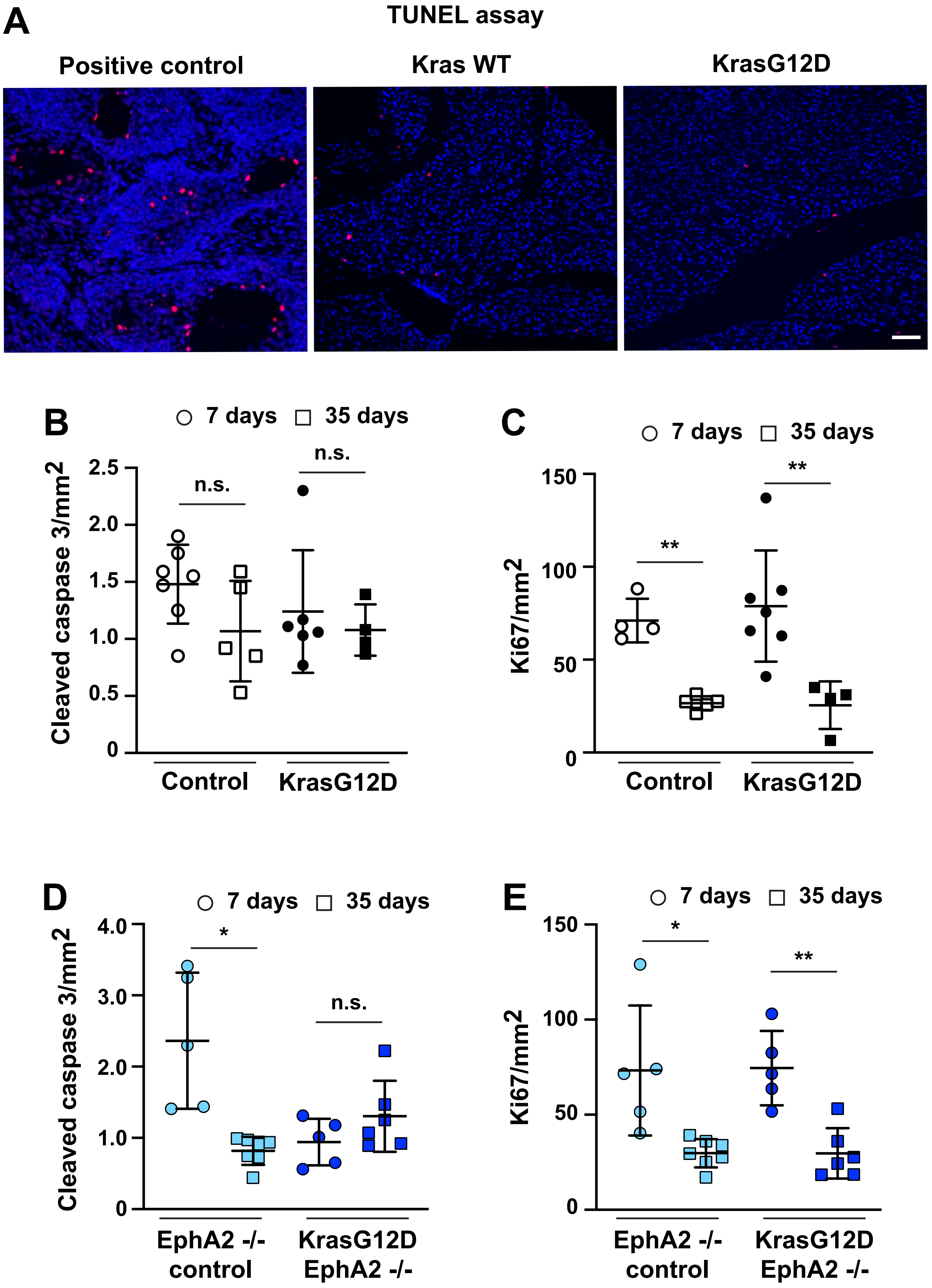


Hill et al
SFig 2

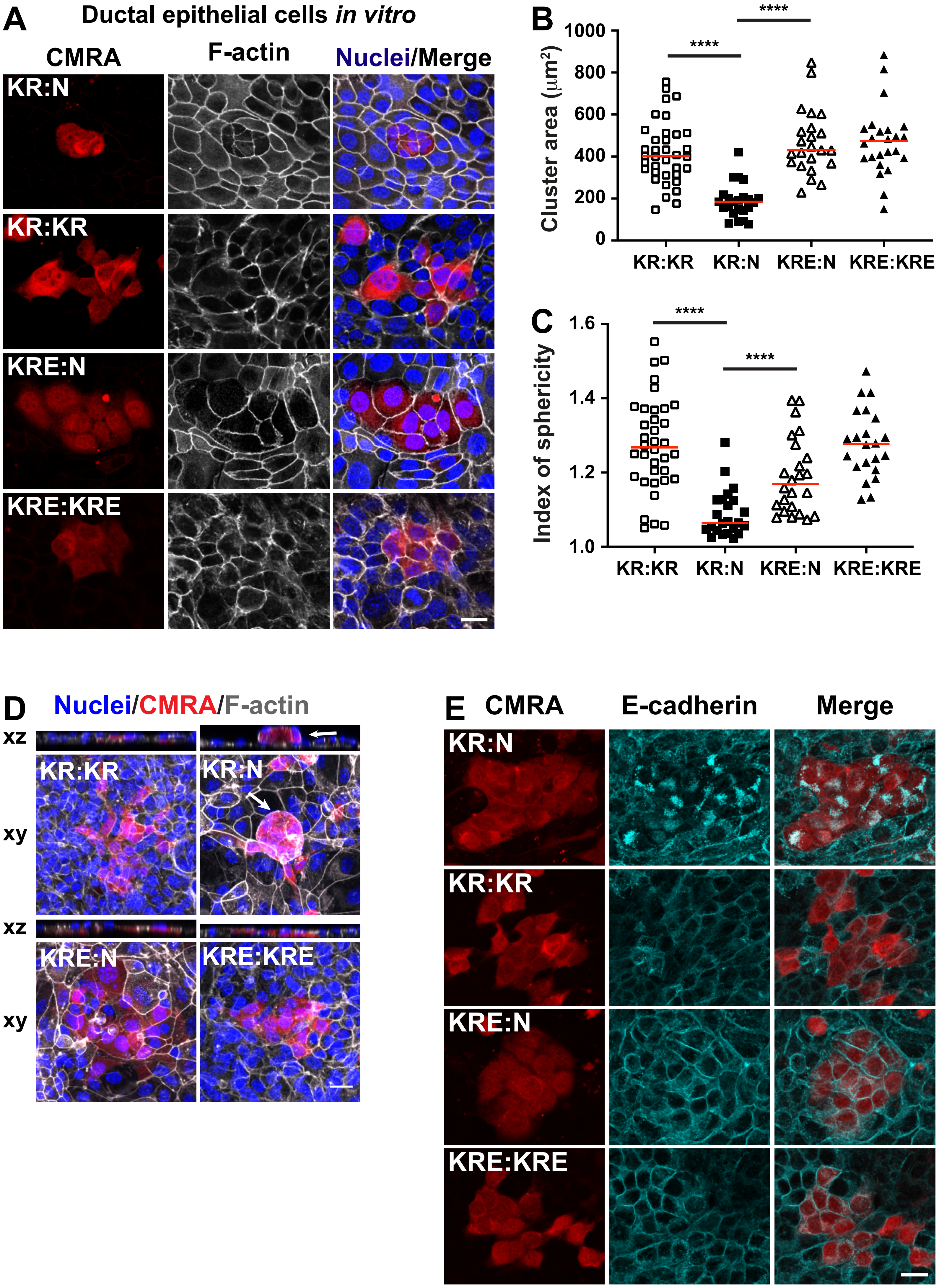


Hill et al
SFig 3

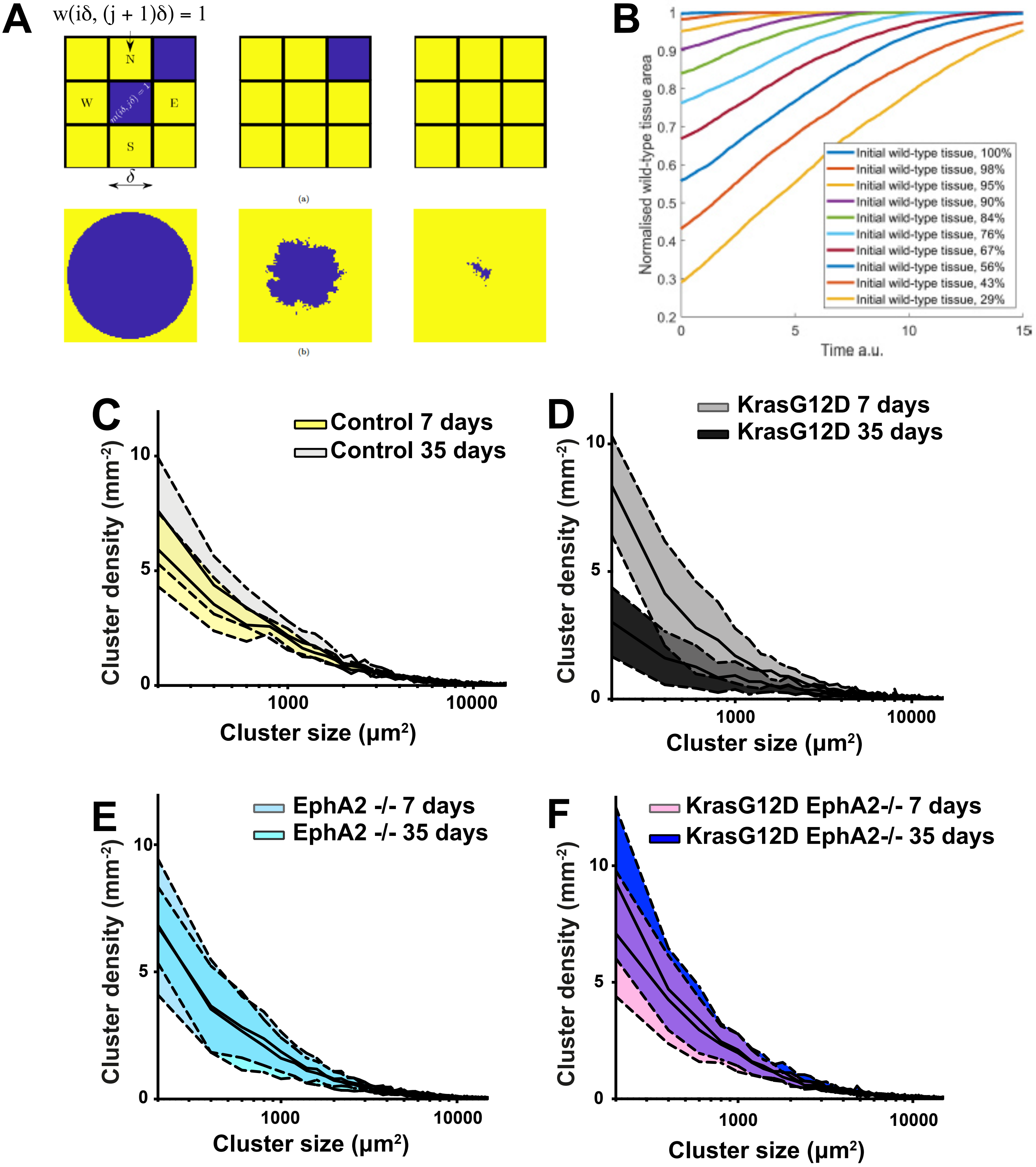


**G**

*p-value*

| Cluster size ($\mu m^2$) | WT v KC | KC v KCE |
|---|---|---|
| 200 | *0.0095* | *0.0094* |
| 400 | *0.0121* | *0.0125* |
| 600 | *0.0067* | *0.0254* |
| 800 | *0.0044* | *0.0057* |
| 1000 | *0.0381* | *0.016* |
| 1200 | *0.01* | *0.0211* |
| 1400 | *0.0194* | *0.0313* |
| 1600 | *0.019* | *0.0047* |
| 1800 | *0.0144* | *0.0133* |
| 2000 | *0.0357* | 0.0823 |
| 2200 | 0.0907 | 0.0603 |
| 2400 | *0.0095* | *0.0303* |
| 2600 | *0.0217* | *0.0168* |
| 2800 | *0.0317* | *0.0318* |
| 3000 | 0.1251 | 0.069 |

**Hill et al**
**SFig 4**
